# Gene body methylation buffers noise in gene expression in plants

**DOI:** 10.1101/2024.07.01.601483

**Authors:** Jakub Zastąpiło, Robyn Emmerson, Liudmila A Mikheeva, Marco Catoni, Ulrike Bechtold, Nicolae Radu Zabet

## Abstract

Non-genetic variability in gene expression is an inevitable consequence of stochastic nature of processes driving transcription and translation. Largely thought to be deleterious to cell fitness, it is not uniform across the transcriptome. This implies the existence of (molecular) determinants affecting the degree of gene expression variability, although this remain poorly understood in multicellular systems. In this study, we found a link between gene body methylation and noise in gene expression in *Arabidopsis thaliana*. More specifically, genes with high levels of noise show low levels of gene body methylation, while genes with lower level of noise in gene expression show higher level of gene body methylation. Most importantly, loss of CpG methylation in gene bodies lead to a significant number of genes displaying higher noise in gene expression. This could be compensated by low but significant gain of non-CpG methylation at promoters of certain genes. Overall, our results show that gene body methylation has a functional role and specifically controls the noise in gene expression for a large number of genes.

## Introduction

Genetically identical organism grown under identical conditions may display different levels of gene expression. This difference or noise in gene expression is ubiquitous across biological systems and may lead to functional differences between individuals (Eldar & Elowitz, 2010). Previous work has shown that transcriptional noise (variability) between genetically identical individuals can arise from the stochastic nature of the molecular processes influencing transcription and translation (Cortijo *et al*, 2019). In animals, gene expression variability is thought to play a role in immune responses (Hagai *et al*, 2018) and, in yeast, it was linked to improved survival under stress (Bishop *et al*, 2007). In plants, gene expression variability is present in the form of seed germination times, where it is a part of a bet-hedging strategy of germination (Abley *et al*, 2024, 2021). More generally, gene expression noise is relatively widespread in plants, and it has been proposed that it could drive developmental patterns and may be important in stress responses (Araújo *et al*, 2017; Cortijo *et al*, 2019; Abley & Locke, 2021). Recent advances in real time monitoring of transcription initiation have shown the extensive contribution of intrinsic noise at the cell level determines tissue level responses (extrinsic noise) especially under environmental stress conditions such as heat and phosphate starvation (Alamos *et al*, 2021; Hani *et al*, 2021). Our understanding of how noise affects inter-plant variability and the mechanisms controlling this noise is limited. A previous study suggested that highly variable genes are smaller, are potentially targeted by a higher number of transcription factors and have a more compacted chromatin environment (Cortijo *et al*, 2019).

Thus, a major source of variation is likely linked to epigenetic changes which are also responsible for different types of cellular memory, including stress memory (Liu *et al*, 2014). Epigenetic changes lead to altered chromatin structure which has been linked to gene expression noise in isogenic cell populations (Wu *et al*, 2017). For example, in Arabidopsis, over-expression of CHR23, a chromatin remodelling ATPase, increased gene expression variability in a distinct subset of genes linked to environmental stress resulting in growth variability (Folta *et al*, 2014). Similar effects have been identified in yeast cells with altered histone deacetylase (HDAC) activity (Weinberger *et al*, 2005).

In *Arabidopsis thaliana*, epigenetic regulation of DNA in form of cytosine methylation is the most common type of DNA methylation (Liang *et al*, 2018). Cytosine methylation can be subdivided based on the sequence context into mCpG (CpG), mCHH (CHH) and mCpGH (CpGH). Gene-body methylation (gbM) consists of an enrichment of CpG methylation within the transcribed regions of a gene, and a depletion at the transcriptional start and termination sites together with an overall depletion of CHG and CHH (where H stands for A,C or T) within the gene (Bewick & Schmitz, 2017). Methylation in CpG, CHG and CHH contexts have different maintenance pathways, but these are not completely independent and there is a link between maintenance of CpG and non-CpG methylation. In particular, CpG methylation at gene bodies triggers accumulation of CHG methylation (Zabet *et al*, 2017b), but (INCREASE IN BONSAI METHYLATION 1) IBM1 demethylase removes this (Saze *et al*, 2008)

Methylation at CHH and CHG context is linked to the transcriptional silencing of transposable elements through the RNA-directed DNA methylation pathway, reviewed by (Matzke *et al*, 2015; Fultz *et al*, 2015), and CHH and CHG methylation in genes and promoter regions, is also linked with gene silencing (Neri *et al*, 2017). Interestingly, the role of CpG methylation at gene bodies remains an open question. It has been speculated that gbM is associated with reduced transcriptional noise or erroneous transcription (Zilberman *et al*, 2008; Huh *et al*, 2013; Neri *et al*, 2017), and recent analysis of single cell RNAseq data of the root quiescent centre cell suggested that gbM is involved in reducing erroneous transcription by reducing intron retention and lowering intrinsic transcription noise compared to unmethylated genes (Horvath *et al*, 2019).

In this study, we investigated the tissue level gene expression noise in *Arabidopsis thaliana* spanning a across a different timescale. More specifically, we identify genes with high expression variability across different timescales and investigated the link between gene expression noise at tissue level and epigenetics (DNA methylation).

## Results

### Gene expression variation in Arabidopsis thaliana

To investigate variation in gene expression, we considered two publicly available timeseries expression datasets produced on leaf 7 of 5-week old Arabidopsis plants: *(i)* the well-watered samples of a progressive drought experiment (drought mock – discovery dataset; (Bechtold *et al*, 2016), which consisted of 14 daily timepoints, and *(ii)* the low light grown samples of a 6 hour high light exposure sampled at 30 minute intervals (high light mock – validation dataset, (Alvarez-Fernandez *et al*, 2021a). Both datasets consisted of four biological replicates at each time point.

First, in order to be able to compute variation in all biological replicates at all time points, we identified a set of probes present in both datasets, which resulted in a total of 19,239 genes for downstream analysis (Supplementary Figure S1). Based on the outcomes of PCA, we removed two discovery and five validation biological replicates from the downstream analysis due to their divergence from the remaining biological replicates (Supplementary Figure S2A-B). The coefficient of variation (CV) was selected as the metric of gene expression variability as previously described (Volfson *et al*, 2006). Based on the distribution of values across the two series (Supplementary Figure S2C), 0.04 was chosen as the cut-off value, to categorise genes as *stable* or *variable* for further analysis. Based on this rule, 1542 genes in the discovery dataset and 1895 in the validation dataset were classed as variable. Of these, 680 were in common between both datasets, which is more than expected by chance (Chi-square test p-value=1.15×10^-6^), and we termed this the consensus list of variable genes (Figure 1A).

**Figure 1.**
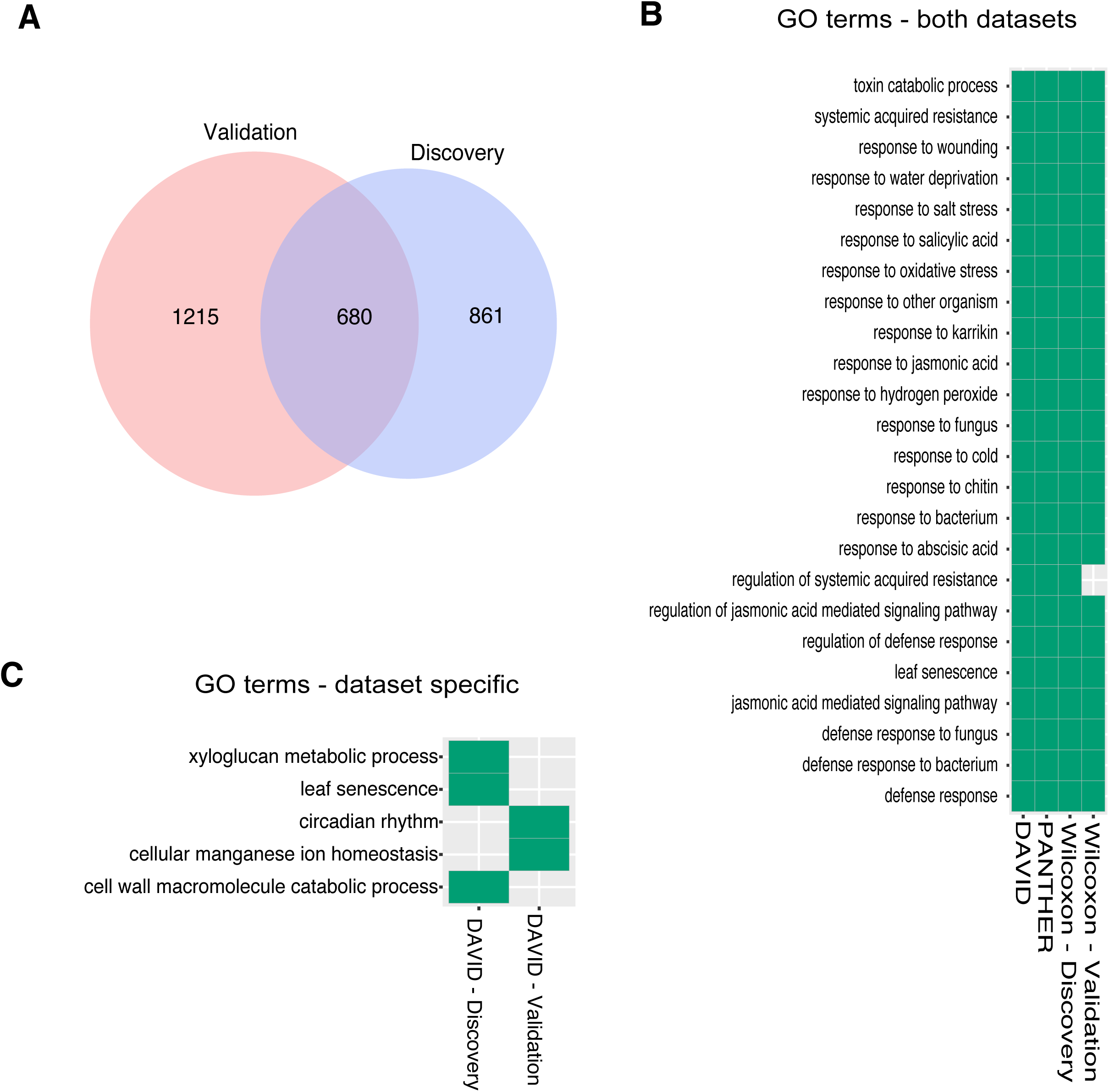
Genes with high variability in gene expression in both datasets. *(A)* Common and unique genes in discovery and validation dataset. The numbers represent the number of genes present in just discovery, just in validation, or both. *(B)* Significantly enriched GO terms (green squares) present in output of DAVID, Panther, and Wilcoxon Rank Sum test in at least one dataset (discovery or validation). *(C)* Same as (B) when considering only genes that are present in either discovery or validation datasets.

GO analysis of the consensus list suggests that most of the genes respond to either an abiotic or biotic stimulus or were involved in the regulation of stress responses (Figure 1B). This suggests that gene expression variability may be involved in stress responses in plants. Interestingly, while validation specific variable genes were enriched for circadian rhythm, the discovery specific genes were enriched in leaf senescence and other processes linked to plant growth and development (Figure 1C). This can be explained by the different timescales (hours vs days/weeks) of both datasets, with the discovery dataset including days/weeks as timepoints and the validation dataset including minutes/hours as timepoints.

### Gene body methylation is associated with stable gene expression

Next, we wanted to investigate if there is a link between variation or stability in gene expression and epigenetic signatures. Thus, we used publicly available bisulfite sequencing data of *A. thaliana* Col-0 ecotype generated in three weeks old leaves by (Stroud *et al*, 2012) and (Stroud *et al*, 2013). The low-resolution profiles analysis of Col-0 BS-seq data showed that the three biological replicates were largely similar with the area of the centromere highly methylated compared to the chromosome arms (Supplementary Figure S3A-C).

To investigate the link between DNA methylation and gene expression, we grouped genes in three classes: *(i)* gene body (gbM) corresponding to only CpG methylation detected in gene bodies; *(ii)* transposable-like corresponding to methylation of at least 5% methylation in CHG and CHH contexts found in gene bodies; and *(iii)* promoter corresponding to methylation in any context identified in the promoter region of a gene. Note that for the latter, we consider the methylation state of the promoter independent of the methylation state of the gene and thus, we could have the case of a promoter being methylated in non-CpG context, but gene body being methylated in CpG context only.

High level of methylation corresponds to lower variation in gene expression for all three categories of genes (Figure 2A, red dots). In contrast, lower level of methylation corresponds to higher variation of gene expression (Figure 2A, blue dots), with few genes displaying both high methylation and high level of variation (purple dots). This is particularly true for gene body methylation in CpG context and promoter methylation in all contexts and these results are reproducible in the validation dataset (Supplementary Figure S4A). Furthermore, the differences in methylation between stable and variable genes in the three groups are statistically significant (except for non-CpG methylation at genes with gene body methylation, Supplementary Figure S5). The statistical analysis further confirm that the most significant difference is observed for gene body methylation (Supplementary Figure S5).

**Figure 2.**
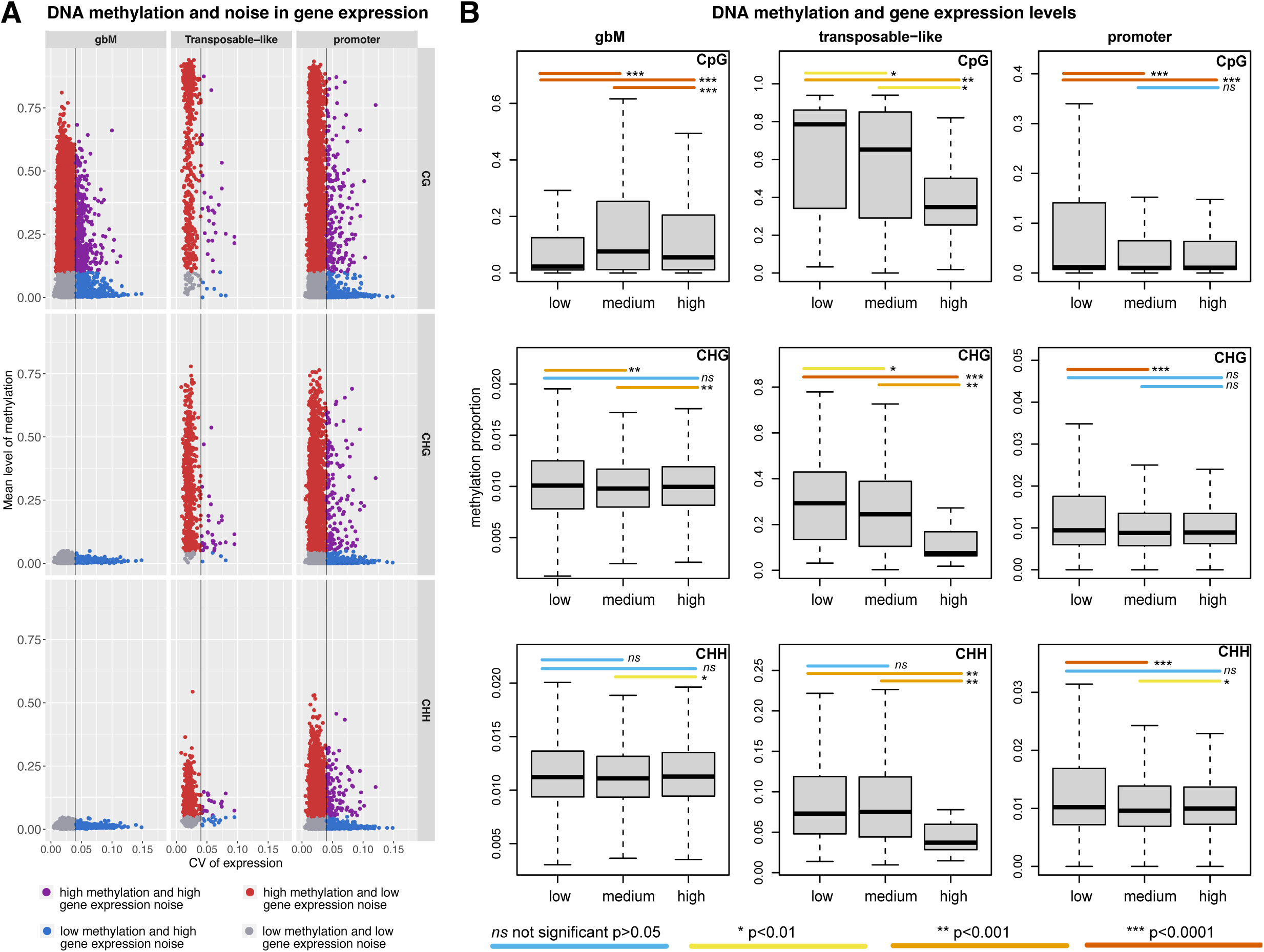
Relationship between methylation level, coefficient of variation for gene expression and level of gene expression. We considered gene expression dataset for controls in (Bechtold *et al*, 2016). *(A)* Link between methylation level and noise in gene expression. We considered separately the case of gene body methylation (gbM), transposon like methylation and methylation at promoters. In addition, we consider context specific methylation, by splitting the methylation in CpG, CHG and CHH contexts. Red points represent genes with high level of methylation and low noise in gene expression, blue points represent genes with low level of methylation and high noise in gene expression, magenta points represent genes with high level of both methylation and noise in gene expression, while grey points represent genes with both low level of methylation and noise in gene expression. (B) Statistical analysis of the relationship between gene expression magnitude and methylation proportion in different methylation contexts and methylation types. The plots are grouped by methylation type: gbM genes (left hand column), genes with transposable element-like methylation (middle column), and promoter methylation (right hand column), divided into different methylation contexts, CpG, CHG, and CHH. In addition, genes are grouped into 3 categories: (i) low with expression values < 8.4; (ii) medium with expression values between 8.4 and 12.05, and (iii) high with expression values > 12.05. Only genes falling into the same category in both discovery and validation datasets were analysed. P-values were calculated using Wilcoxon rank-sum test. The colour of the lines between two boxes indicates the statistical relationship between the two distributions as calculated by Wilcoxon rank-sum test as indicated on the figure.

Considering that DNA methylation is associated to transcriptional silencing, one possibility is that methylated genes are less expressed, and low expression could explain the lower variability. To explore this possibility, we evaluated the correlation between methylation levels and levels of expression. For genes with TE like methylation, this was indeed the case, with highly expressed genes having low or no methylation (Figure 2B and Supplementary Figure S6). Nevertheless, for gene body methylation and promoter methylation, high levels of methylation were not associated with a low level of expression (Figure 2B and Supplementary Figure S6). In fact, medium and high level of gene body methylation correspond to higher level of gene expression compared to low levels of gene body methylation (Figure 2B). Again, these results are reproducible in the validation dataset confirming that they are not affected by the dataset used in our analysis (Supplementary Figure S4A). Given that the validation and discovery datasets are measured at different timescales (hours vs weeks), this suggests that the results are independent of effects induced by developmental or circadian regulation.

### Loss of gene body methylation in MET1 mutants increases gene expression variability

So far, our results have shown a correlation between gene body methylation and stability of gene expression. To further investigate if there is a functional link between the two, we considered data in MET1 mutants that are hypomethylated in CpG methylation. Investigating the MET1 mutants allows us to focus on whether the CpG methylation is controlling the stability in gene expression. We used methylation and expression datasets from two MET1 mutants: *(i) met1-1* that results in a loss of approximately three quarters of CpG methylation and *(ii) met1-3* that results in a complete loss of CpG methylation (Supplementary Figure S7). The CpG methylation that is retained in *met1-1* mutant is mainly located in promoters, TEs and non-CpGs methylated regions, but almost all gene body methylation is lost (Supplementary Figure S8A) (Catoni *et al*, 2017). This contrasts with *met1-3* mutant where virtually all CpG methylation is lost, including in promoters, gene bodies and TEs.

In both *met1-1* and *met1-3* mutants, there is a subset of genes that are upregulated compared to Col-0, but also a large subset of genes that maintain same level of expression (Supplementary Figure S8B-C). To investigate the effect of methylation loss on gene expression variability, we considered only genes (and promoters) that lost CpG methylation but maintained the same level of gene expression compared to Col-0. This ensures that the effects on variability of gene expression are not caused by the change in levels of gene expression, but only by the loss of CpG methylation. Independent of where the methylation is located (gene bodies, promoters) or the type of methylation (transposon like), a significant higher number of genes show increased variability in expression in *met1-1* mutant (Figure 3). This is statistically significantly higher than the number of genes with decreased variability in gene expression. The largest effect is observed for gene body methylation, where more than 4000 genes increase the variability in gene expression upon loss of CpG methylation compared to less than 2000 genes that display a decrease in variability in gene expression. Figure 4A-C shows example of three genes (AT4G12040, AT1G30320 and AT5G27450) that completely lose gene body methylation in *met1-1* mutant and show an increase in the C.V. In contrast, AT5G64460 retains approximately half of gene body methylation in *met1-1* mutant and no corresponding change in C.V. (Figure 4D).

**Figure 3.**
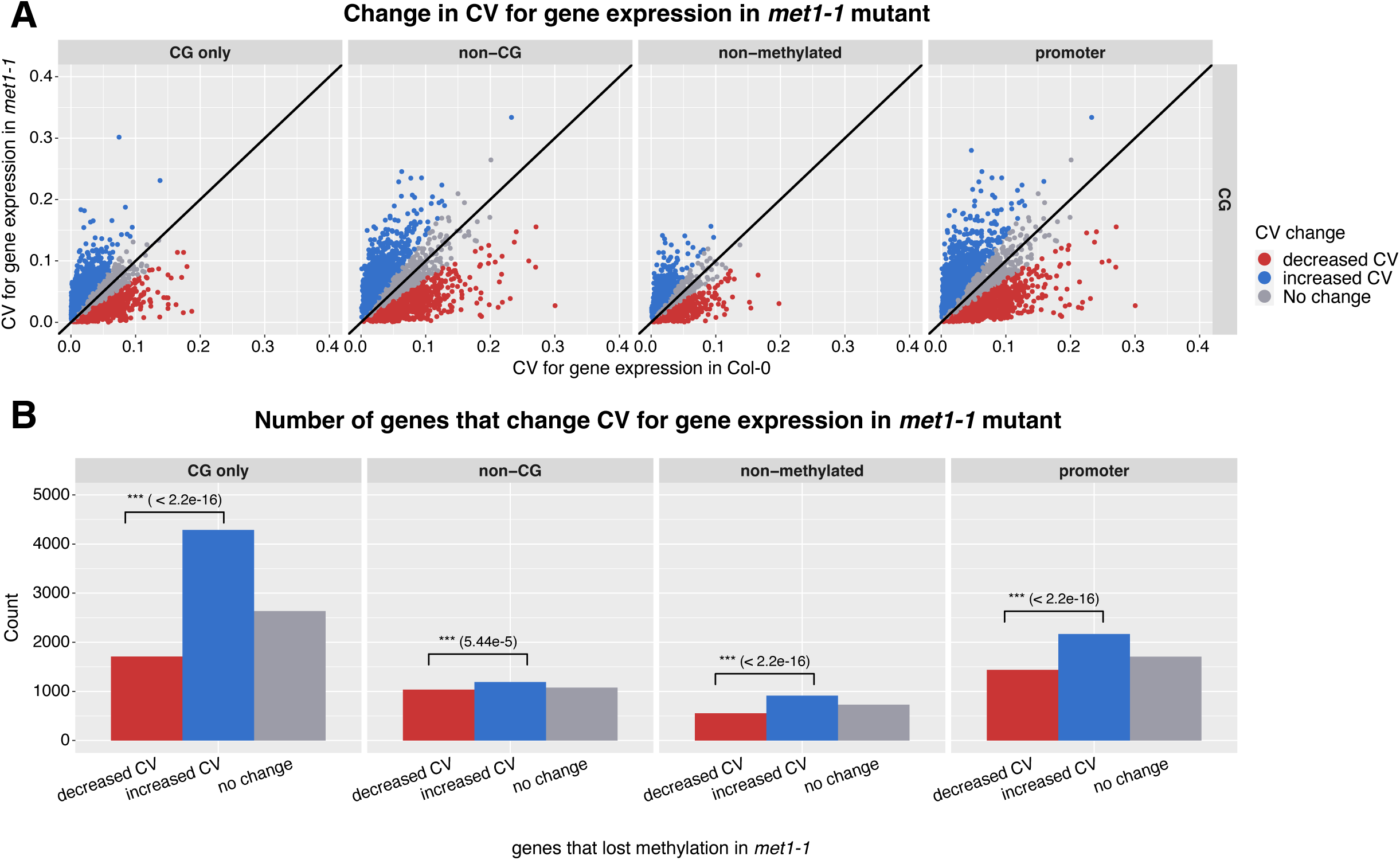
Changes in noise in gene expression in met1-1 mutant. *(A)* Comparison between the Coefficient of Variation (CV) in WT and *met1-1* plants; data from (Catoni *et al*, 2017). We considered in this analysis only genes without significant change in gene expression between WT and mutant but show loss of DNA methylation (overlap with a hypomethylated DMR in *met1-1*). Genes are grouped based on their coefficient of variation change: (blue) increased for genes with CV fold change greater than 0.5, (red) decreased for genes with CV fold change less than –0.5, and (grey) no change for those in-between. (B) Number of genes in each group shown in panel A.

**Figure 4.**
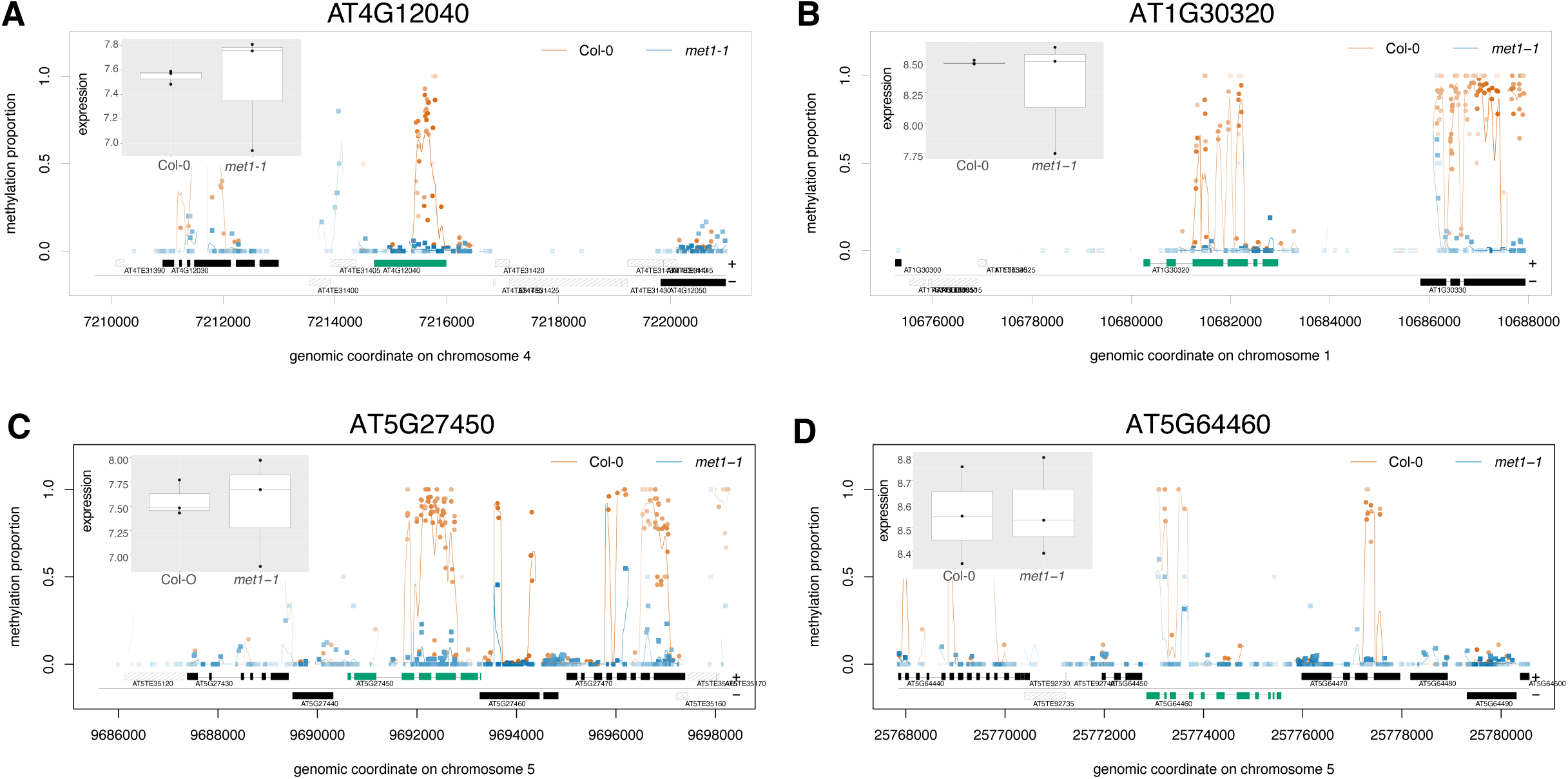
Loss of gene body methylation results in increase in noise in gene expression. We considered four examples: *(A)* AT4G12040, *(B)* AT1G30320, *(C)* AT5G27450 and *(D)* AT5G64460. The red colour represents methylation in *met1-1* mutant and the blue colour in Col-0 plants. The gene of interest is highlighted by marking the exons with green. The inset represents the expression levels for the corresponding gene. In (A-C) the complete removal of gene body methylation results in a large increase in gene expression noise. In (D), the retention of approximately half of gene body methylation results in maintenance of the same level of gene expression noise.

Contrary to what we have seen in *met1-1* mutant, in *met1-3* we observed similar number of genes that increase or decrease the variability in gene expression, indicating that although both mutants affect the C.V. of many genes, only in *met1-1* this results in an overall increase of variation in expression (Supplementary Figure S9A-B). There are several possibilities that could explain these results. First, the set of genes that lost DNA methylation in *met1-1* could be more prone to display an increase in noise upon loss of DNA methylation. However, the large majority of genes that have lost CpG methylation are common to both *met1-3* and *met1-1* mutant. This suggests that the effect in C.V. variation observed in *met1-1* is not driven by specific set of genes with altered methylation specifically in the *met1-1* mutant.

Secondly, it is possible that *met1-3* mutant, being a complete lack of function allele, accumulates larger epigenetic and transcriptomic changes and, consequently, more indirect effects occur. To investigate how many of these changes in variability in gene expression in *met1-3* are also found in *met1-1*, we investigated the overlap of genes in the two mutants that display either increase or decrease in variability of gene expression (Supplementary Figure S10). The largest overlap is for genes that lose gene body methylation in both mutants and increase variability of expression (1433 genes), which is approximately three times higher than genes that lose gene body methylation and decrease variability in gene expression (576 genes).

### Gene expression variability in epigenetic recombinant inbred lines

Furthermore, to investigate if the asymmetric change in C.V. observed in *met1-1* was masked in *met1-3* background due to the high number of genes that change expression, we took advantage of epigenetic recombinant inbred lines (epiRILs) generated by a cross between Col-0 and *met1-3* plants, and propagated by single seed descend (Catoni & Cortijo; Reinders *et al*, 2009). Each epiRIL line plant displays parts of the genome inherited from a Col-0 epigenetic state and parts from the genome in *met1-3* epigenetic state, but they all contain a wild type allele of MET1 gene. By analysing their WGBS dataset, we were able to annotate in each line the parts of the genome that are Col-0 or *met1-3*. We then performed RNA-seq for ten epiRILs line in two biological replicates. In each epiRIL line, genes were labelled as either Col-0 or *met1-3*, depending on whether the genomic region where the gene was located were Col-0 or *met1-3*. We selected the genes that appeared at least two times in Col-0 background and two times in *met1-3* background in the ten epiRIL lines and grouped their expression levels based on the background they come from. This resulted in generating pseudo Col-0 and pseudo *met1-3* replicates for each gene (see *Materials and Methods*). Similar as we did previously, we selected the genes that maintained expression level independent on whether they were in Col-0 or *met1-3* background and estimated the C.V. in gene expression among the pseudo replicates in Col-0 and *met1-3*. Our results showed that only 624 genes with gene body methylation increased variability in expression when in *met1-3* background compared to Col-0, which was higher than the number of genes that decreased variability (567), although not statistically significant; see Supplementary Figure S11. Similar results were also observed when the methylation was in the promoter of the gene. This indicates that similar numbers of genes increasing or decreasing variation in *met1-3* mutant can also be observed in epiRILs, and therefore is not a direct consequence of the *met1-3* mutant allele or an indirect effect of the strong transcriptional changes occurring in this mutant background.

### Asymmetry of CpG methylation in met1-1 mutant is linked with variability in gene expression

Since approximately one quarter of CpG methylation is retained in *met1-1*, one possibility is that methylation is lost preferentially at genes that increase the variability in expression, which appears not to be the case (Supplementary Figure S12A). Alternatively, the *met1-1* specific increase in variation could reflect variation at level of the DNA molecules analysed in the sampled tissue, at level of alleles or single cell epigenetic landscape. This could be explained by a partial or incomplete activity of MET1, which is consistent with the lower stability of MET1 protein observed in this mutant (Catoni et al 2017).

MET1 methyltransferase is responsible for maintaining methylation at symmetric CpG sites and, thus, the observation of asymmetry in CpG could be associated with incomplete MET1 activity. Therefore, we look specifically if methylation could be maintained at both cytosines in a CpG site or that one cytosine loses the methylation while the other cytosine maintains it. Our results show that at genes that increase variability in expression in *met1-1* mutant, there is a significant decrease in methylation symmetry at CpG sites that are still methylated (Supplementary Figure S12B). Nevertheless, this asymmetry in methylation at CpG sites is also observed at genes that display a lower variability in gene expression, suggesting that asymmetry of DNA methylation alone cannot explain the buffering in noise in gene expression observed in *met1-1* mutant.

### Decreased variability of expression in met1-3 mutant correlates with a reduced acquisition of non-CpG methylation at promoters of a subset of genes

From the transcription data we have analysed, and also from previous work, we observed increased non-CG methylation occurring in the *met1-3* mutant, which is likely the consequence of incorrect splicing of the histone demethylase IBM1 responsible for removing non-CpG methylation in gene bodies (Inagaki *et al*, 2017; Rigal *et al*, 2012; Zabet *et al*, 2017a) Click or tap here to enter text.. However, when we looked at genes for which variability in expression was affected in the *met1-3* mutant, we noticed that non-CpG (both CHG and CHH) hypermethylation is more pronounced in promoters of genes with a decreased C.V. (Figure 5A-B), while this is not evident in wild type or the *met1-1* mutant (Figure S13). This effect is also absent in gene bodies, which display a similar increase in non-CpG methylation irrespective of the change in C.V. (Figure 5C-D). It is worthwhile noting that this low but significant accumulation of non-CpG methylation at promoters of genes does not lead to changes in gene expression at the genes considered in our analysis, i.e. we only selected genes that are not differentially expressed in *met1-3* for this analysis (see Figure S8C). Considering that repeated DNA has been associated with the ability to attract methylation (Catoni et al, 2017), we investigated the presence of repetitive elements near (within 10Kb) promoters of genes that increase or decrease variability of expression in *met1-3* mutant. We found an average of one annotated transposable element (TE) in the proximity of promoters of all group of genes (Figure S14A), however, we observed that there is preferential association of certain families near genes that decrease variability of expression (Figure S14B), including ATGP (5 and 6), ATCOPIA (21, 4, 43, 62 and 12), VANDAL (2, 3, 5, 7 and 21) and ATHILA4. Furthermore, other TE families are preferentially located near genes that increase variability of expression (Figure S14C), including COPIA (10, 14, 18A, 19, 31, 32, 39, 46, 48, 53, 63, 73, 77 and 86) and VANDAL (10, 18 and 20). These results suggest that CHH and CHG methylation accumulate in the *met1-3* mutant at specific TE families, which could partially compensate the variability of expression gained for the loss of CpG methylation in bodies of downstream genes. Therefore, the effect of increased variability in expression found in the *met1-1* mutant, at least for some genes, could be buffered in *met1-3* by the deposition of non-CpG methylation. While our results demonstrate a relationship between DNA methylation and variability of gene expression, it is however likely that multiple mechanisms are involved in controlling the variability of expression, which could be directly or indirectly linked to epigenetic regulation.

**Figure 5.**
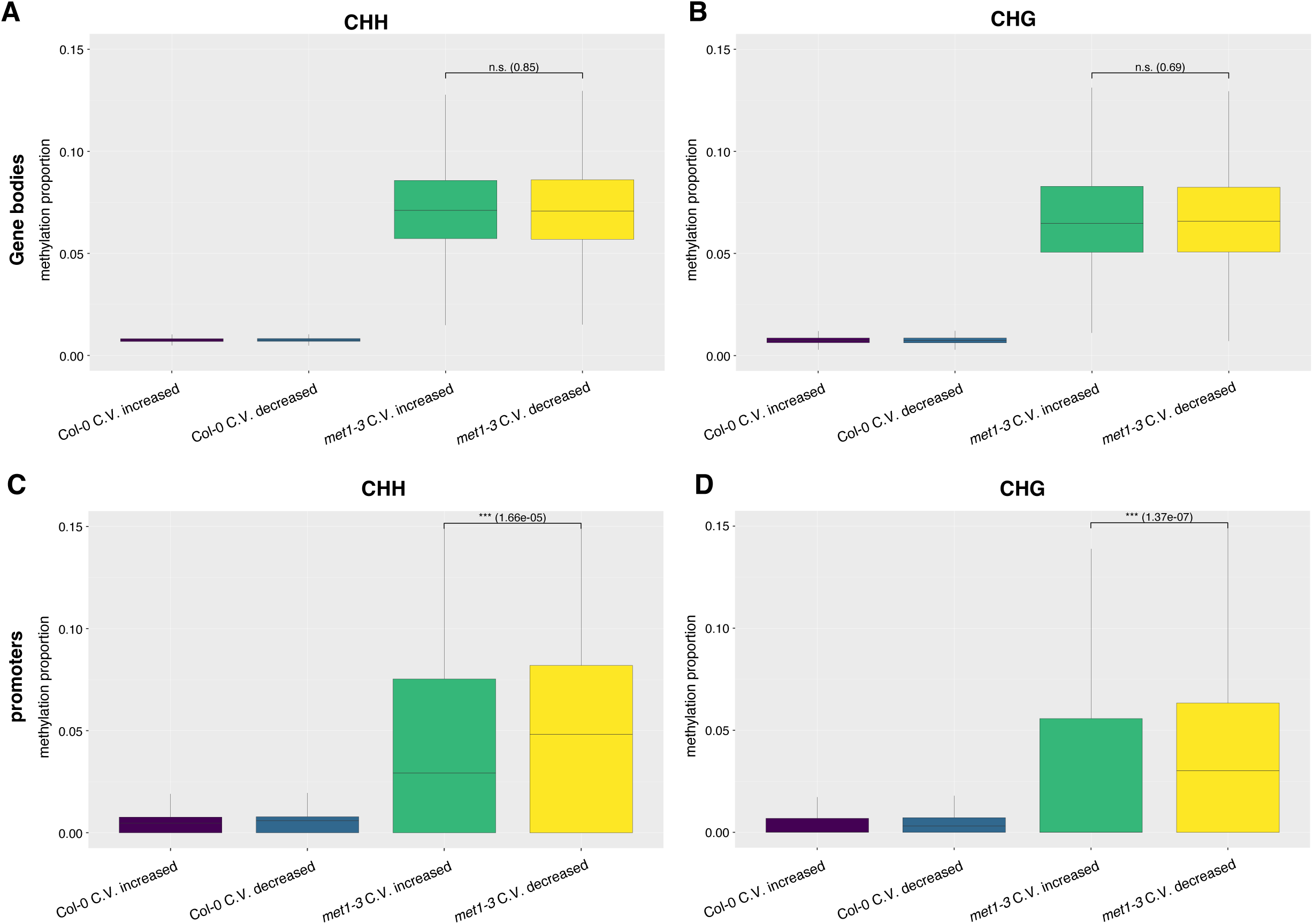
non-CpG methylation in the met1-3 mutant at genes that display increase or decrease of noise in transcription. Methylation level of cytosines in (*A* and *C*) CHH and (*B* and *D*) CHG context located (*A-B*) within genes or (*C-D*) promoters of genes that displayed an increase or decrease in variability in gene expression in *met1-3* while maintaining the same expression levels as in Col-0. We performed the Mann–Whitney U test with corresponding P-value added to the plots.

## Discussion

Noise is hardwired in biochemical systems, either as an intrinsic component from biochemical reactions or as extrinsic factor that is propagated through regulatory pathways (Spudich & Koshland, 1976; Arkin *et al*, 1998; Elowitz *et al*, 2002). Previous work has identified some potential sources of variation in gene expression between isogenic populations, but the finer details and the mechanism responsible for regulation of transcription variation are currently not fully understood. Our previous work showed that a reduction in noise in gene expression without slowing down the response time of genes can be achieved by: *(i)* increasing metabolic cost associated with the gene (Zabet & Chu, 2010), *(ii)* having a negative feedback loop (the gene encodes for a TF that represses its own expression)(Zabet, 2011) or *(iii)* the TF regulating the activity of the gene performing facilitated diffusion (a combination of one-dimensional scan and 3D diffusion) (Schoech & Zabet, 2014). Here we characterised genes that display noise in gene expression in a single leaf of the model plant *A. thaliana*, and specifically investigated the role of DNA methylation in the regulation of gene expression noise.

Our results show that in two different timeseries datasets consisting of 13 evenly spaced timepoints collected across different timescales, the genes displaying high variability in expression are enriched in genes related to stress response. This is consistent with previous research, which has identified that genes responsible for stress response are highly variable in yeast (Bar-Even *et al*, 2006), and in Arabidopsis, across a 24h diurnal period, with different classes observed between the day and night time (Cortijo *et al*, 2019). Both our datasets contained samples taken exclusively during the light period, but interestingly, the discovery dataset was collected over a 14 a day period indicating that the link between gene expression variability and stress response is independent of timescales and the developmental stage of the rosette leaf. It has been suggested, however, that developmental stochasticity is caused by noisy gene expression, which may vary between tissues and/or cell types (Araújo *et al*, 2017). Both datasets were taken from the fully expanded rosette leaf 7, comprising of a number of tissues and cell types, representing transcriptional heterogeneity in heterogenous cell types, and therefore may have a less significant effect on mean expression levels of the entire tissue.

Next, we performed a systematic analysis and found a link between DNA methylation in gene bodies and variability in gene expression. In particular, we found that genes with gene body methylation (DNA methylation only in CpG context located in gene bodies) displayed low noise in gene expression, while genes lacking gene body methylation displayed higher noise in gene expression. This link is not restricted to gene body methylation, but is also present at promoters, where methylated promoters displayed stable gene expression while unmethylated promoters were affected by noise in gene expression. We identified this association using bulk expression datasets in both the 6h and 14 days datasets. Recently, another study using scRNA-seq datasets identified a similar anticorrelation between gene body methylation and noise in gene expression (Horvath *et al*, 2019).

One possibility is that there is only a correlation between gene body methylation and noise in gene expression and that there is no functional relationship between the two. To further dissect if this link between DNA methylation and noise in gene expression, we analysed expression and methylation datasets in two MET1 mutants, *met1-1* and *met1-3*. In particular, *met1-1* results in loss of approximately three quarters of CpG methylation genome wide, while *met1-3* in complete loss of CpG methylation (Stroud *et al*, 2013; Catoni *et al*, 2017). Our results show that the loss of gene body methylation in *met1-1* resulted in increase in variability of gene expression (three time more genes increase the noise in gene expression than genes that reduced it). This is statistically significant, and it is true when we select only genes that maintain the same level of gene expression, excluding that this is linked to genes changing expression levels. Furthermore, we also found that there are more genes that display higher noise even upon loss of CpG methylation at promoters. Interestingly, this was not the case with *met1-3* mutant, where there is no predominant increase in noise in gene expression.

Interestingly, it seems that low but significant accumulation of non-CpG methylation in promoters can explain the decrease in transcription variability at genes in *met1-3* (Figure 5). Overall, we have a model where CpG methylation buffers noise in gene expression and loss of CpG methylation corresponds to an increase in transcription variability, but this can be compensated by an increase in non-CpG methylation in promoters of certain genes.

In *met1-1*, there is loss of CpG methylation mainly in gene bodies, but in *met1-3*, there is loss of CpG methylation genome wide, including repeats and transposable elements. One possibility is that the loss of CpG methylation at TEs can increase non-CpG methylation at promoters of genes, by RNA-dependant DNA methylation (RdDM) mechanisms (Erdmann & Picard, 2020). Our results showed that the promoters of both genes that display an increase or a decrease in variability in *met1-3* have nearby TEs, but different families of TEs are present at promoters of genes that decrease variability of expression compared to genes that increase variability of expression in *met1-3*.

Finally, DNA methylation has been proposed to be involved in repression of gene transcription. In plants, while this seems to be the case at genes with transposon like methylation (having both CpG and non-CpG methylation), it is not the case for genes having only gene body methylation, which display medium and high expression levels (Lister *et al*, 2008). Given the metabolic cost associated with establishment and maintenance of DNA methylation, this raises the question of what the functional role of DNA methylation in gene regulation is. Here, we show that DNA methylation, especially in gene bodies, acts as a buffer to the biochemically hardwired noise in gene expression. Similar role is also observed with DNA methylation at promoters, albeit, with a weaker effect of DNA methylation on the noise in gene expression. It is worthwhile noting that here we focused on gene transcription, which does not always accurately predict concentration of protein (Gygi *et al*, 1999). As such, it is possible that post-transcriptional mechanisms would moderate the effects of variation in mRNA concentrations.

## Materials and Methods

### Datasets

We used two publicly available microarray mock expression data: *(i)* mock drought series (Bechtold *et al*, 2016) (GSE65046) with four biological replicates at each time point and measurements performed every day for 14 days (resulting in 56 samples) and *(ii)* mock data from high light series (Alvarez-Fernandez *et al*, 2021b) (GSE78251) with four biological replicates at each time point and measurements performed every 30 minutes for 6 hours (52 samples). In both series, tissue from *Arabidopsis thaliana* leaf 7 of a separate plant for each biological replicate was harvested, and sequenced using CATMA microarrays (Sclep *et al*, 2007). The plants in both series were 5 weeks old at the start of measurement series.

The two datasets used CATMA microarray version, and a consensus probe-set was constructed, featuring only probes present in both (Bolstad *et al*, 2003) datasets (Supplementary Figure S1). The joined datasets were normalized together, using cyclic loess normalization method (Bolstad *et al*, 2003).

We also used gene expression (3 biological replicates) in *met1-1* and *met1-3* mutants and their corresponding WT plants from (Catoni *et al*, 2017). The data was further processed using Robust Multi-Array Average expression algorithm, without normalisation component, which was carried out separately, using *normalize.ExpressionSet* function of the affyPLM (Brettschneider *et al*, 2007), with loess method and antilog transformation. A linear model fit was processed to compute Bayesian statistics for the comparison using eBayes function. p-values were FDR corrected using *p.adjust* function of stats package. Mean, standard deviation, and coefficient of variation of expression were calculated between biological replicates for each gene separately for each expression series.

For DNA methylation, we used publicly available bisulfite-converted sequencing data for 3-week-old leaves harvested from three wild type Col-0 ecotype (Stroud *et al*, 2013, 2012). In addition, we used Bisulfite-converted sequencing data for 2-week-old leaves harvested from wild type Col-0 Arabidopsis, *met1-1*, and *met1-3* (Catoni *et al*, 2017). The data was processed following protocol outlined by (Catoni *et al*, 2018). EpiRILs lines has been obtained by Reinders et al (2009), and RNA-seq and WGBS were generated as part of this study.

### Genomic data analysis

Sequenced reads have been processed with Trimmomatic (Bolger *et al*, 2014) to remove adapters and discard low-quality reads. The cleaned reads were mapped to Arabidopsis reference genome (TAIR10 version) using TopHat2 (Kim *et al*, 2013) for RNAseq and Bismark (Krueger & Andrews, 2011) for WGBS datasets.

For RNAseq, Picard tool (available from http://broadinstitute.github.io/picard/) was used to discard duplicated reads and mapped reads from TopHat2 were subsequently counted using htseq-count (Anders *et al*, 2015). Then, the raw count per gene was used to determine differentially expressed genes between tested condition using DESeq2 (Anders & Huber, 2010) with default parameters. Genes with a significance p-value below 0.05 after Benjamini and Hochberg correction were considered to be differentially expressed.

For WGBS, mapped reads have been de-duplicated with Bismark and the cytosine report has been produced for each sample analysed. To account for non-converted DNA, we applied a correction according to (Catoni and Zabet 2021). Briefly, the number of methylated reads were decreased as: m*= max(0, m – nc) (where m* is the corrected number of methylated reads, m is the raw number of methylated reads, n is the total number of reads and c is the conversion rate)."

To determine parent chromosomal recombination in epiRILs, we call methylation in each gene using *analyseReadsInsideRegionsForCondition* function in DMRcaller package (Catoni *et al*, 2018). Then, we filtered genes that contain only CpG methylation (mCpG > 50%; CHG/CHH < 1%; min cytosine covered > 5), producing a list of 20,520 epigenetic markers. Then using the same function, we calculated the CpG methylation level of these markers for all epiRILs analysed, relative to the methylation level calculated in the wild-type sample. We when used the R *smooth.spline* function (spar = 0.6) to smooth data across chromosomes and at any chromosomal location we assigned “Col-0-like” methylation to smoothed values above 0.8 and “met1-like” methylation for values below 0.2, leaving the rest classified as epi-heterozygous or undefined (in average less than 5% of each epiRIL genome).

### Identification of Differently Methylated Regions

To identify differently methylated regions (DMRs) we used the DMRcaller package (Catoni *et al*, 2017). In particular, we used *bins* method with the statistical test “score”. Moreover, for CpG methylation, the minimum proportion difference was set to 0.4 (40% difference), while for CHG and CHH, to 0.1 (10% difference), in accordance with lower magnitude of methylation in those contexts. DMRs were assigned to genes with minimum overlap size set to 50. Genes were split into three groups – those overlapping only hypomethylated DMRs, those overlapping only hypermethylated DMRs, and those overlapping both hypomethylated and hypermethylated DMRs both. This analysis was performed for all three methylation contexts and was repeated for promoters.

### Classification of genes by methylation

Gene methylation data was split into three categories, based on their methylation in WT: *(i)* genes with transposable element-like methylation (CHG or CHH context methylation proportion higher than 0.05), *(ii)* genes with gene-body methylation (CpG context methylation proportion higher than 0.1 and CHG and CHH lower than 0.05) and *(iii)* non-methylated genes. We selected for our analysis genes were the number of total reads in WT or MET1 mutants was at least 25. This was done for each biological replicate separately, and for both gene-bodies and promoters. In MET1 mutants, genes hypermethylated in CpG context, were discarded.

### Gene expression comparison between WT and MET1 mutants

Genes with significant expression change, were defined here as FDR higher than 0.05, and log2 of expression fold change higher than 1 or lower than -1. They were excluded from further analysis, in order to avoid bias caused by expression difference. The list of genes and the change in gene expression variability and methylation in *met1-1* mutant are included in Supplementary Table S1 and Supplementary Table S2, while the list of genes in *met1-3* mutant are included in Supplementary Table S3 and Supplementary Table S4

For the epiRILs, we assigned each gene as either being in Col-0 or *met1-3* epigenetic profile in each of the ten epiRIL lines analysed, based on the genomic analysis described above (also see Supplementary Table S5). Genes for which an epigenetic profile was not clearly identified (either because they were close to a recombination point or in a epi-heterozygous area), genes that had mean FPKM value lower than 0.5 (for epiRILs and WT) and genes that had more than one replicate with expression of 0 (for epiRIL and WT) were removed from the analysis. We then selected the genes that are at least in two epiRIL lines in Col-0 epigenetic profile and in two epiRIL lines in *met1-3* epigenetic profile and grouped their expression levels based on the assigned genotype, considering them “pseudo-replicates”, pseudo Col-0 or pseudo *met1-3*. Each epiRIL line had two biological replicates and each gene can have different number of pseudo replicates varying between four and sixteen since the epiRILs contain different recombination points between the original Col-0 and *met1-3* chromosomes used in the original cross. We selected randomly three raw counts values for Col-0 and three for *met1-3* and used DESeq2 to call differential gene expression and used an FDR threshold of 0.05, and no value for log2 of expression fold change. This was repeated 200 times and selected the genes that were not differentially expressed in any of the 200 iterations. Genes that were not differentially expressed were taken forward and we computed the C.V. of the expression in pseudo Col-0 or pseudo *met1-3*. Next, we performed 200 iterations sampling three FPKM values for Col-0 or for *met1-3* and computed the C.V. We then reported the mean C.V. from the 200 iterations as the C.V. of that gene in the corresponding background. Genes that display a difference in C.V. between Col-0 and met1-3 pseudo replicates of at least 0.5 were considered to increased/decreased the variability in gene expression.

The list of genes and the change in gene expression variability and methylation in epiRIL lines are included in Supplementary Table S6 and Supplementary Table S7.

### Gene Ontology analysis

We used two external tools to perform GO enrichment analysis: *(i)* Panther Overrepresentation Test (Mi *et al*, 2009) and *(ii)* DAVID (Huang *et al*, 2009). In addition, we also used *go_enrich* function of the GOfuncR (Maintainer & Grote, 2024) package, using the Wilcoxon rank-sum test method and GO term annotation for Arabidopsis genes from (Berardini *et al*, 2004). For Panther, the output was specified to feature False Discovery Rate for correction, and Fisher’s exact test type. Panther and DAVID analyses of the consensus geneset were conducted twice, using two backgrounds – first, the “all Arabidopsis genes” background, built into both tools, and, second, the custom background, generated from all genes that were included into the Genomic Ranges object. All other Panther and DAVID analyses made use of the custom background, and no background was required for Wilcoxon analysis. The results of enrichment analysis were compared using q-value metrics. Go_enrich were corrected to match the output of Panther and DAVID manually, using p.adjust function of the stats package.

### Data access

All WGBS and RNA-seq data sets from this study have been submitted to the NCBI Gene Expression Omnibus (GEO; http://www.ncbi.nlm.nih.gov/geo/) under accession numbers GSE255301 (RNA-seq) and GSE255302 (WGBS). All scripts used for pre-processing and further analysis can be accessed at https://github.com/ZastaJak/athaGE_variation_and_buffering.

---

**Supplementary Figure S1.**
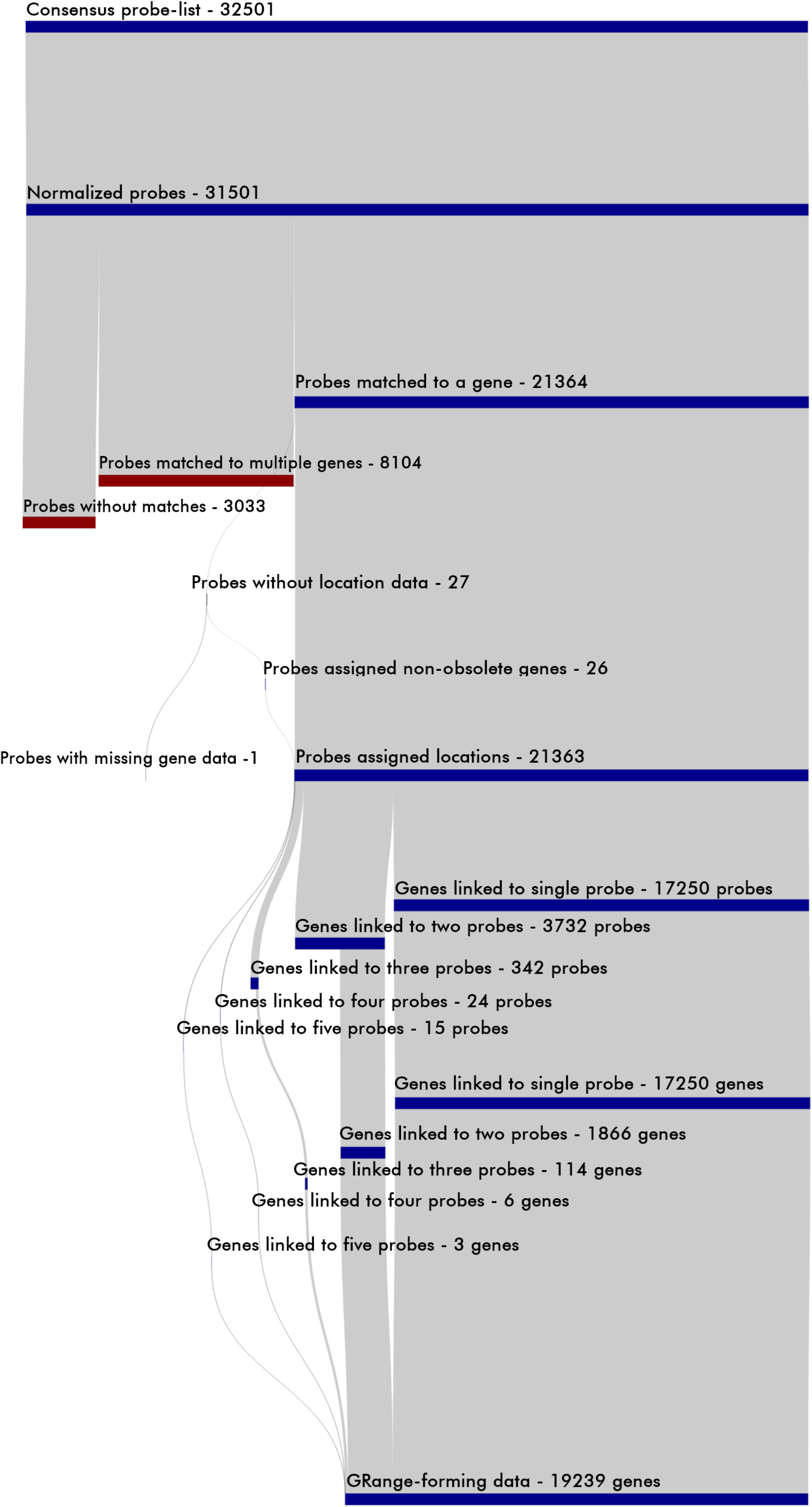
Microarray expression data processing. Sankey diagram representing the steps taken to process the merged discovery and validation microarray data. Bars in red represent rejected probes, bars in blue represent probes or genes passed on to further analysis. Each flow represents the number of genes/ probes passed for the next analysis step.

**Supplementary Figure S2.**
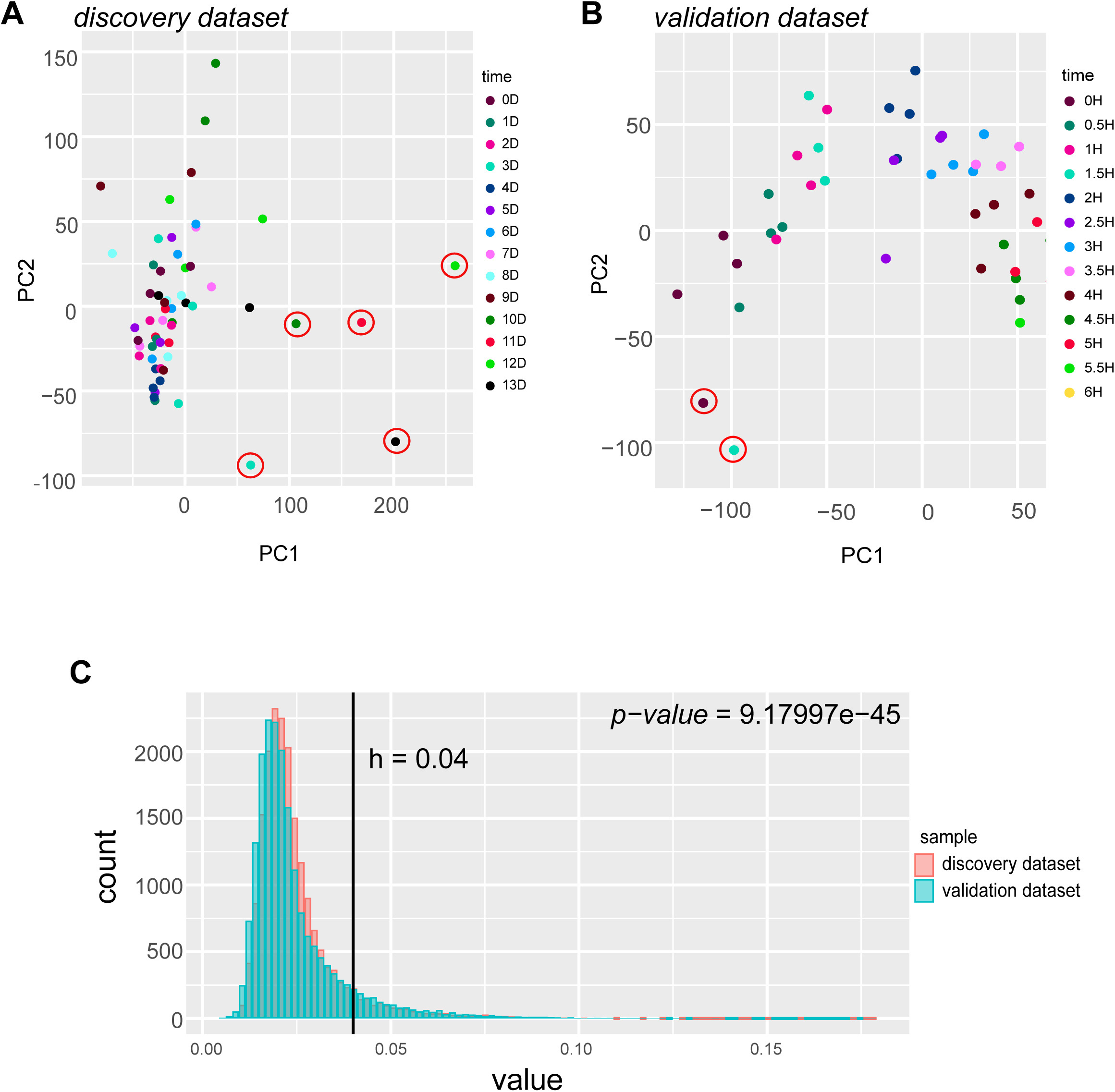
Principal Component Analysis (PCA) and analysis of variance. *(A)* Discovery dataset PCA (Bechtold *et al*, 2016). Each datapoint corresponds to a single biological replicate and biological replicates are grouped by colour and are labelled according to their time point in hours. Datapoints highlighted in red circles were removed from further analysis. *(B)* Validation dataset (Alvarez-Fernandez et al., 2021) PCA. *(C)* Coefficient of variation for discovery (red) and validation (blue) datasets. The vertical line represents a cut-off point set at 0.04 based on the distribution of the values across the two-time series. Values above this cut-off point were classified as variable.

**Supplementary Figure S3.**
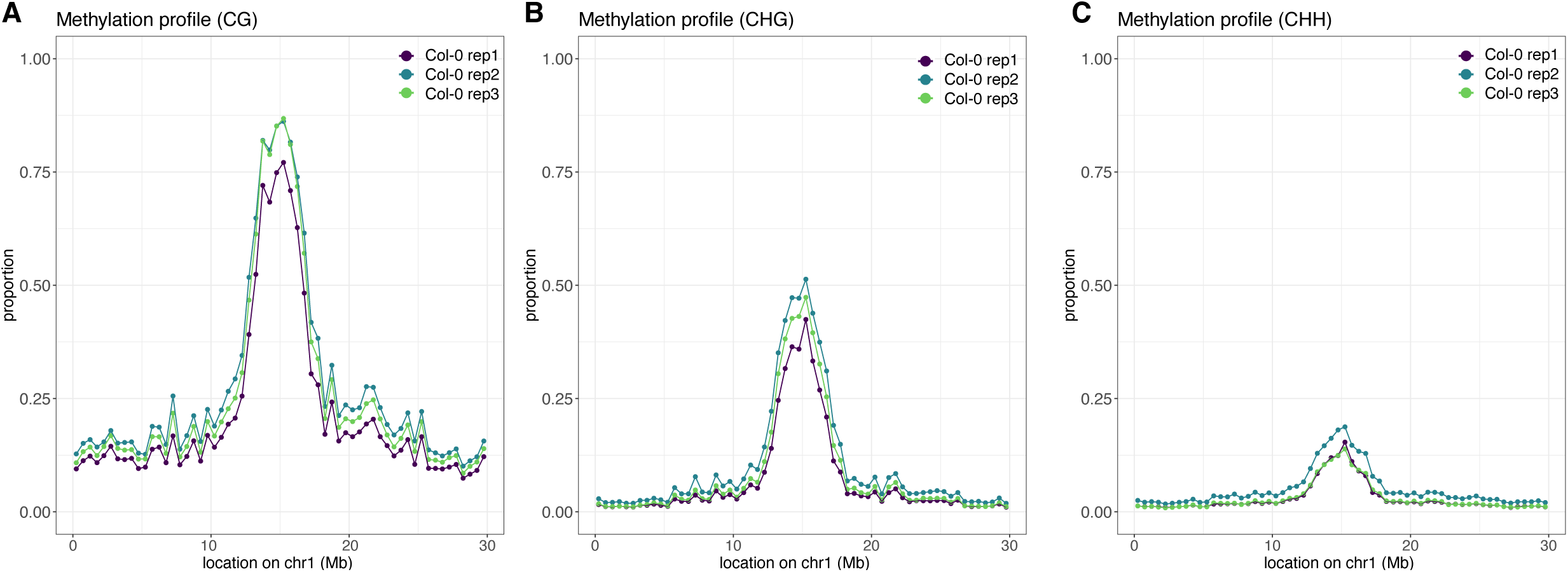
Low resolution (500000 bp) methylation profiles of chromosome 1 in Col-0. *(A)* CpG, *(B)* CHG and *(C)* CHH context.

**Supplementary Figure S4.**
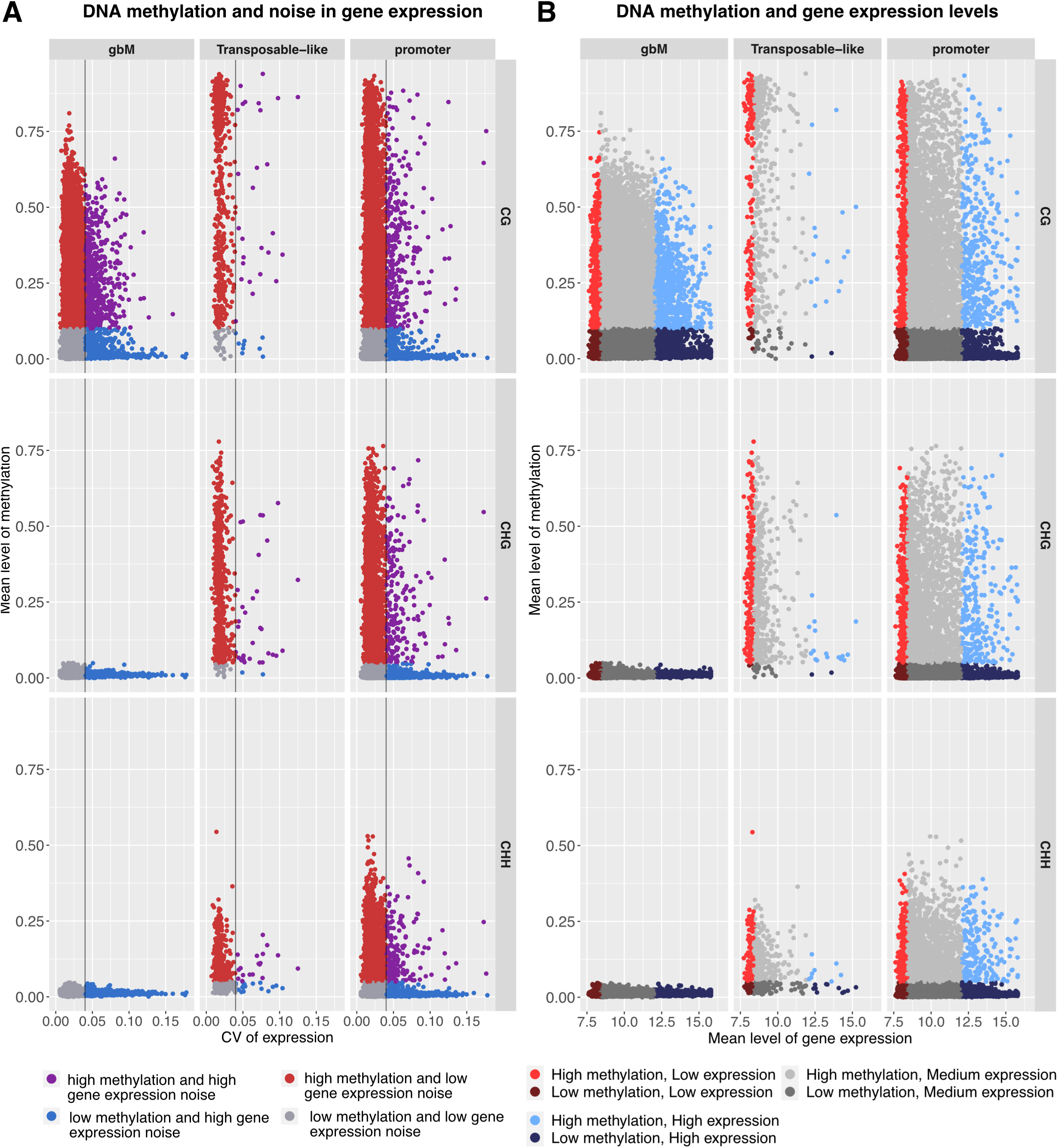
Relationship between methylation level, coefficient of variation for gene expression and level of gene expression in a different dataset. Same as Figure 2 in main manuscript but using the control datasets from (Alvarez-Fernandez *et al*, 2021b).

**Supplementary Figure S5.**
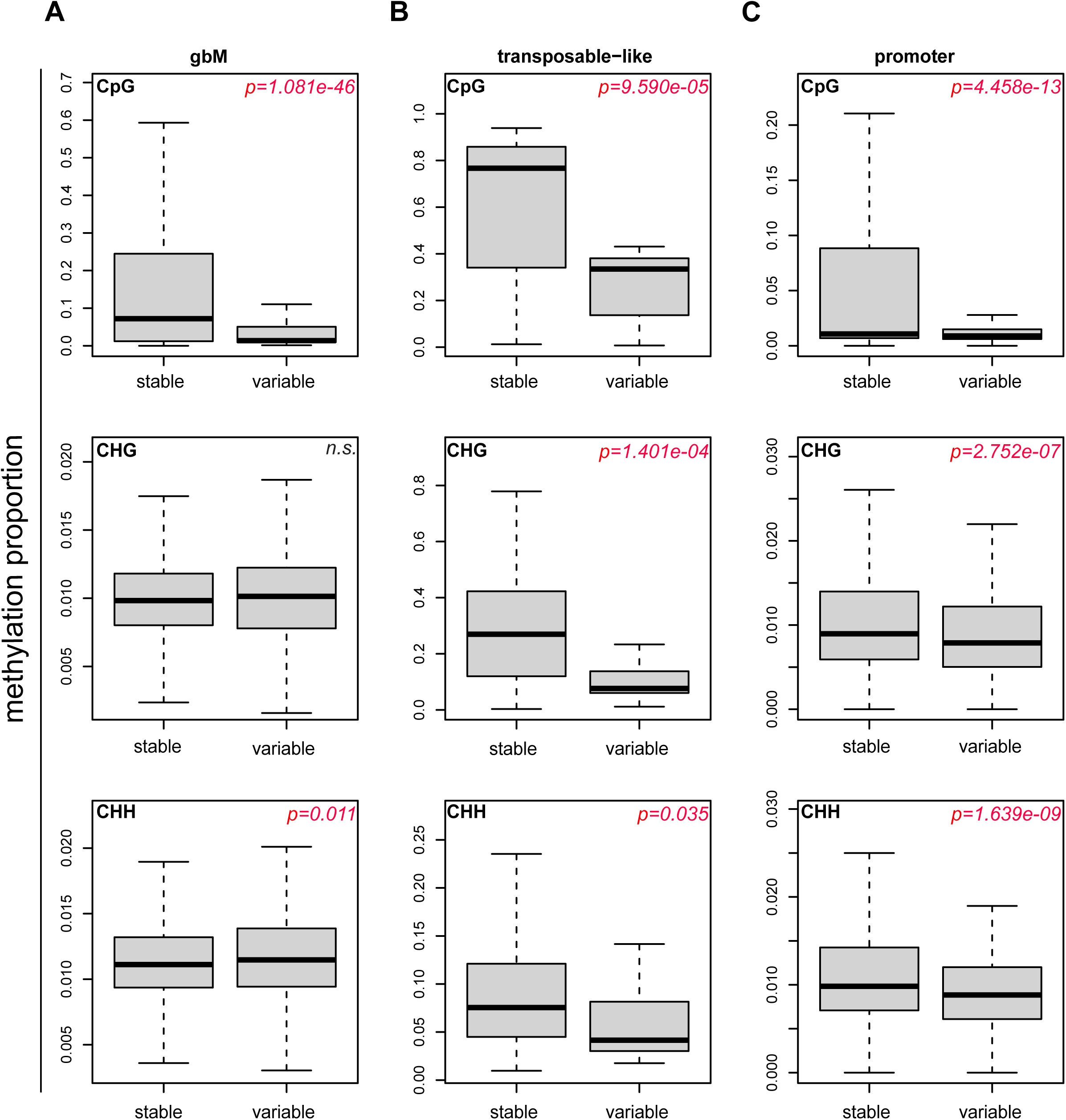
Relationship between CV of expression and methylation proportion in different methylation contexts. Genes were divided into two categories based on the covariance of expression: variable (CV > 0.04) and stable (CV < 0.04). We considered separately the case of: *(A)* gene body methylation (gbM), *(B)* transposable-like elements, *and (C)* promoters. P-values were calculated using the Wilcoxon rank-sum test. Significant p-values are provided, *n.s.* – not significant.

**Supplementary Figure S6.**
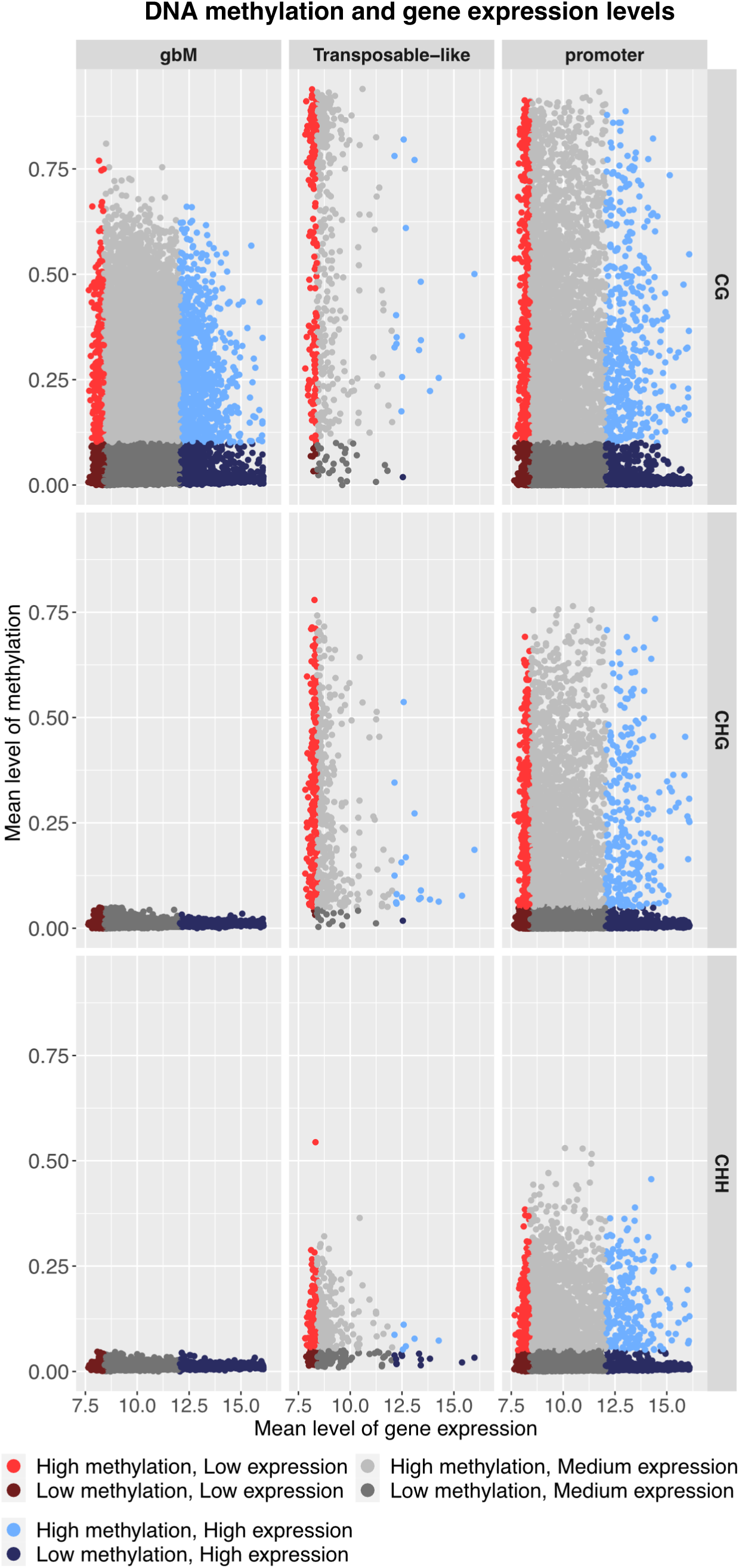
Link between methylation level and level of gene expression. We considered separately the case of gene body methylation (gbM), transposon like methylation and methylation at promoters. In addition, we consider context specific methylation, by splitting the methylation in CpG, CHG and CHH contexts. Red points represent genes with high level of methylation and low noise in gene expression, blue points represent genes with low level of methylation and high noise in gene expression, magenta points represent genes with high level of both methylation and noise in gene expression, while grey points represent genes with both low level of methylation and noise in gene expression.

**Supplementary Figure S7.**
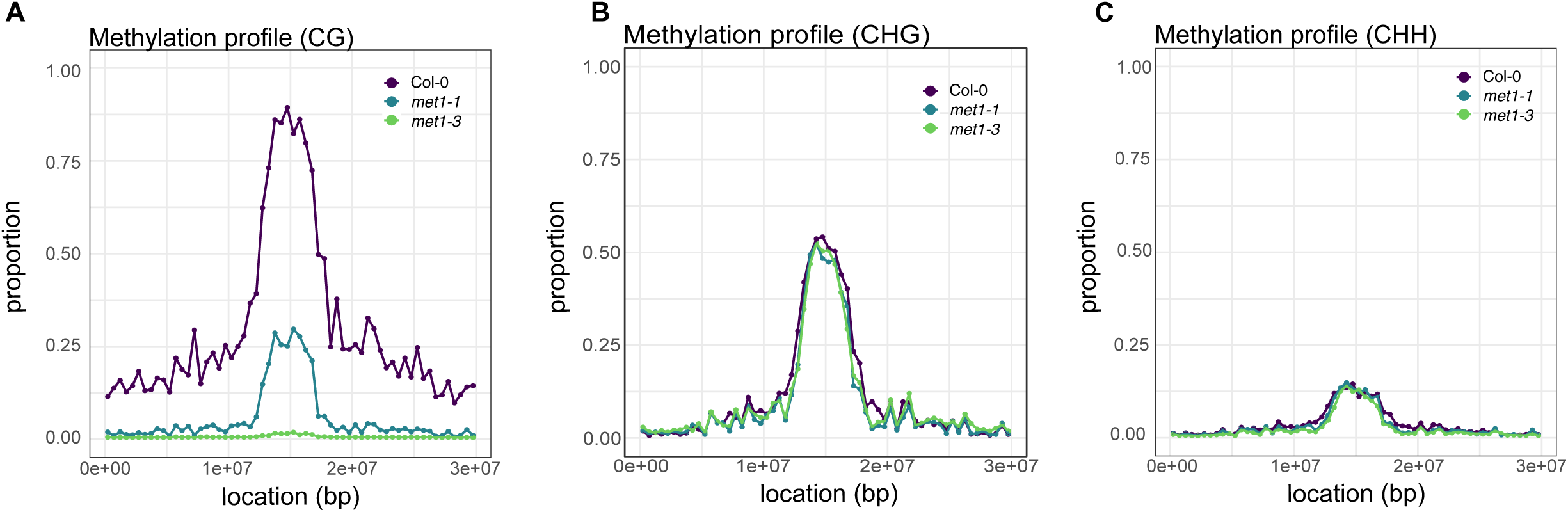
Low resolution (500000 bp) methylation profiles of chromosome 1 in met1-1 and met1-3 mutants. We considered the methylation contexts individually: *(A)* CpG, *(B)* CHG and *(C)* CHH. Black line (Col-0), blue line (*met1-1)* and green line (*met1-3)*.

**Supplementary Figure S8.**
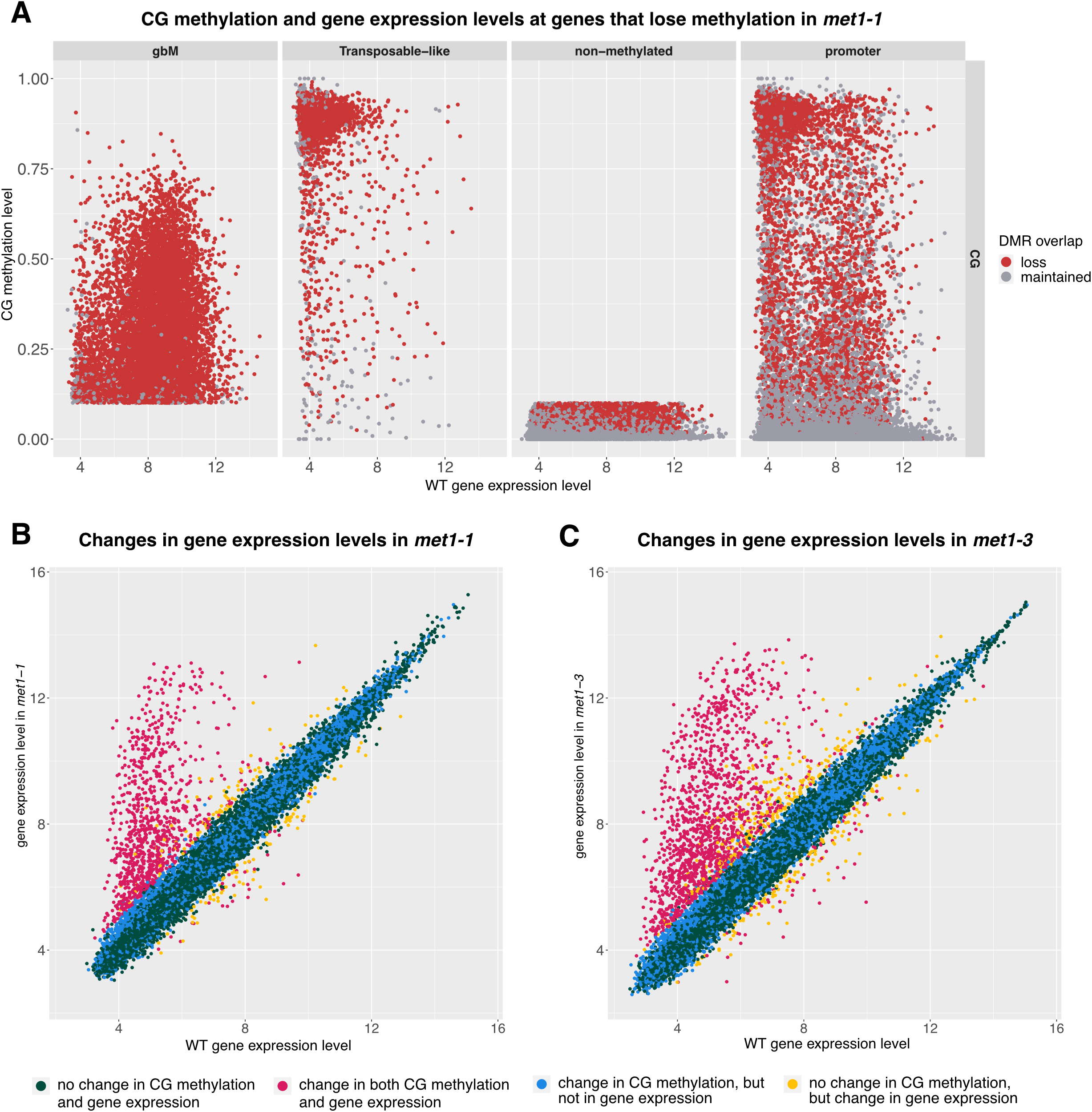
Changes in gene expression in MET1 mutants. *(A)* We plot the methylation level and gene expression level for all genes in WT plants; data from (Zabet *et al*, 2017b). Genes that lose methylation in *met1-1* mutant are highlighted with red. We considered separately the case of gene body methylation (CpG methylation proportion higher than 0.1 but CHG and CHH methylation proportion equal to or below 0.05), transposon like methylation (CHG or CHH methylation proportion higher than 0.05), no methylation (CpG methylation proportion below or equal to 0.1, and CHG and CHH below or equal to 0.05) and methylation at promoters (CpG methylation proportion above 0.1 or CHG or CHH above 0.05). In addition, we consider context specific methylation, by splitting the methylation in CpG, CHG and CHH contexts. *(B-C)* Comparison between gene expression levels in *(B)* WT and *met1-1* mutant, and *(C)* WT and *met1-3* mutant. Genes are separated into four groups: no change in methylation and gene expression levels (green); no change in DNA methylation but change in gene expression (yellow), loss of DNA methylation but no change in gene expression (blue); and change in both DNA methylation and gene expression (red).

**Supplementary Figure S9.**
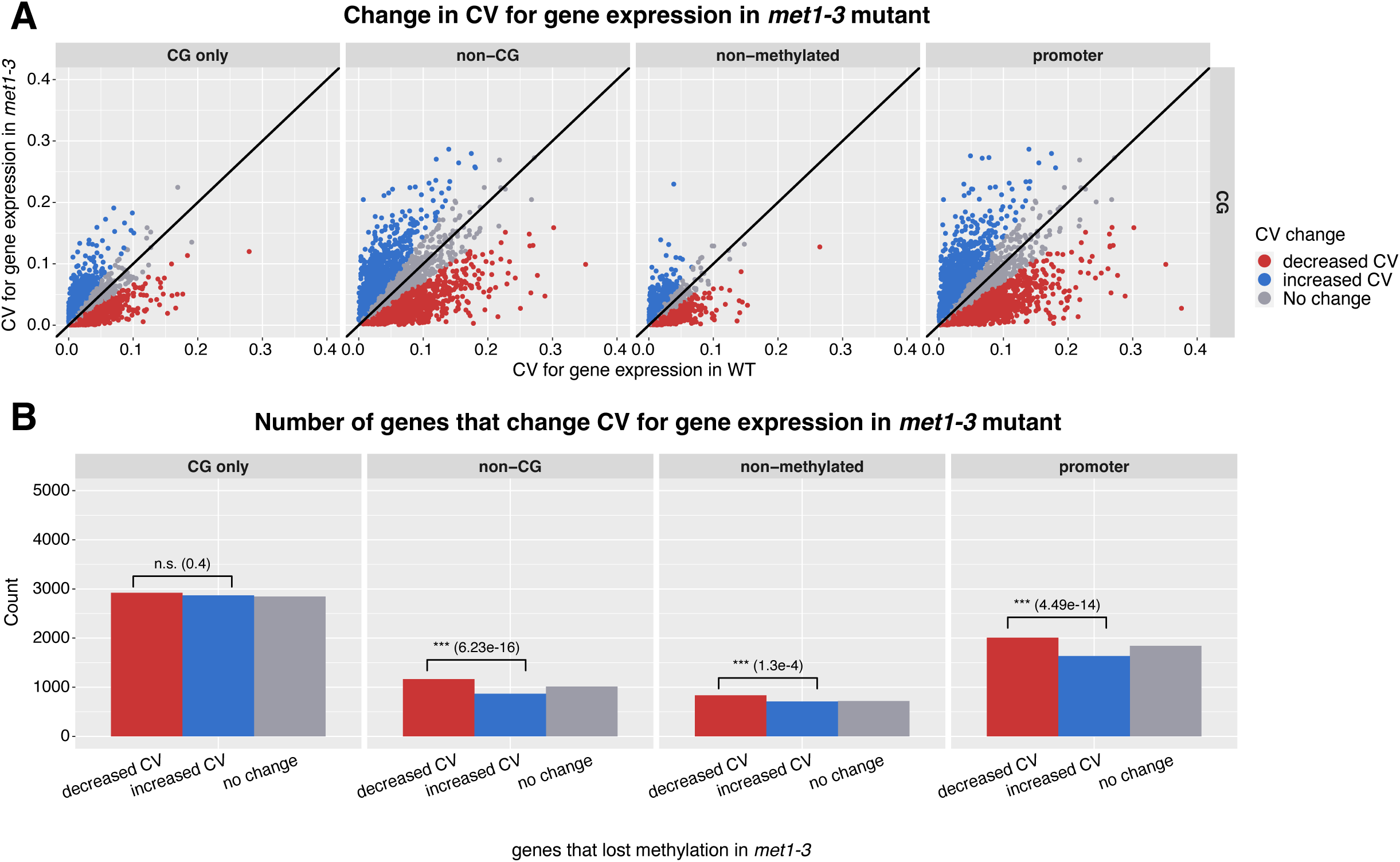
Changes in noise in gene expression in met1-3 mutant. *(A)* Comparison between the Coefficient of Variation (CV) in Col-0 and *met1-3* plants; data from (Catoni et al., 2017). We considered in this analysis only genes without significant change in gene expression between WT and the mutant but displaying loss of DNA methylation (overlap with a hypomethylated DMR in *met1-3)*. Genes are grouped based on their coefficient of variation change: (blue) increased for genes with CV fold change greater than 0.5, (red) decreased for genes with CV fold change less than –0.5, and (grey) no change for those in-between. *(B)* Number of non-differentially expressed genes with loss of DNA methylation between WT and *met1-3* in the different categories shown in panel A.

**Supplementary Figure S10.**
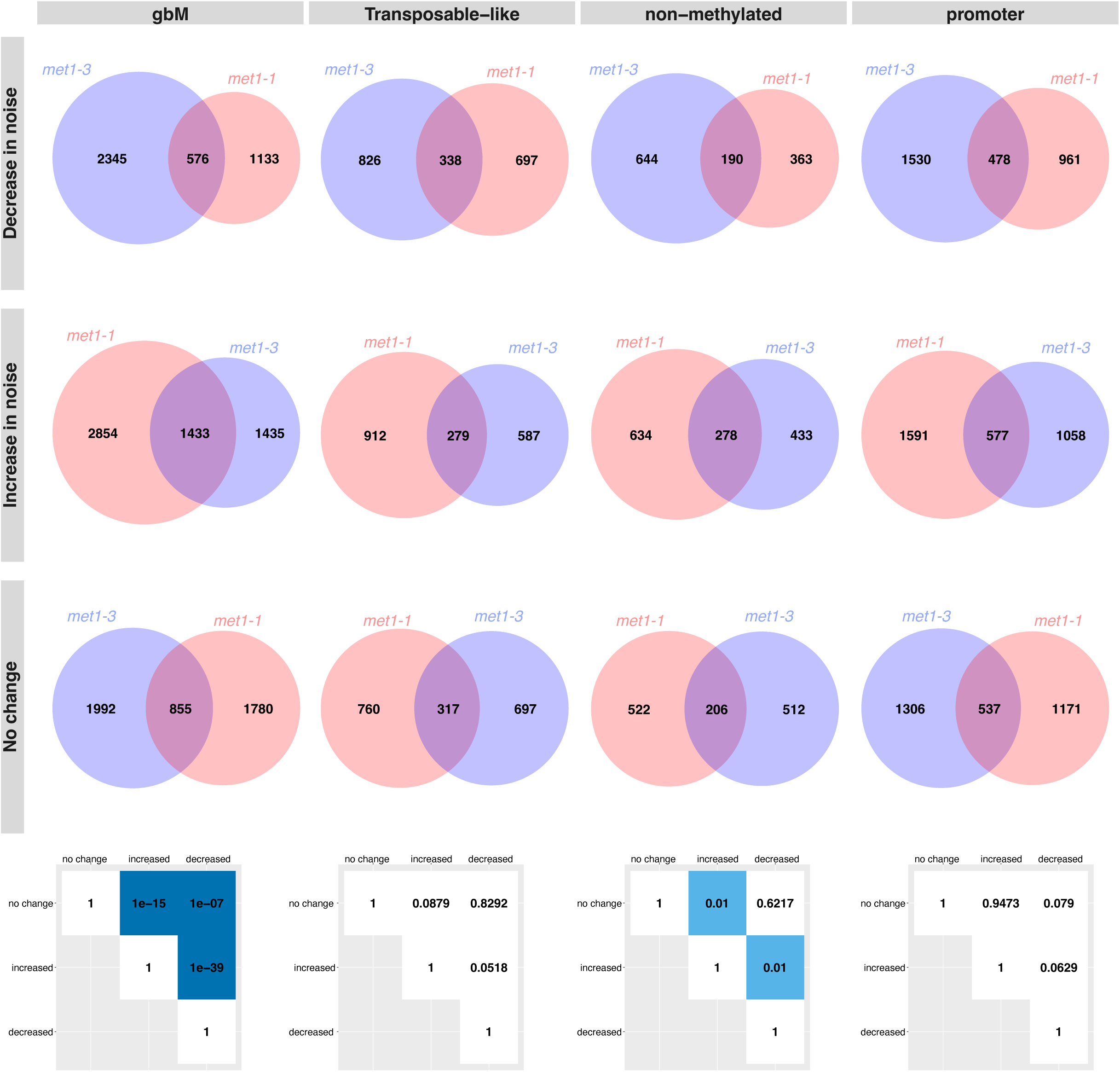
Comparison between changes in noise in gene expression in MET1 mutants. *(A)* Venn diagrams comparing the number of gene that decrease, increase or do not change the coefficient of variation in *met1-1* (red) and *met1-3* (blue) mutants. Genes are split into four categories: *(i)* gene body methylated genes, *(ii)* transposon like methylated genes, *(iii)* un methylated genes and *(iv)* promoter methylated genes. *(B)* Statistical test for the overlaps for each category in panel (A), calculated using Fisher’s Exact Test.

**Supplementary Figure S11.**
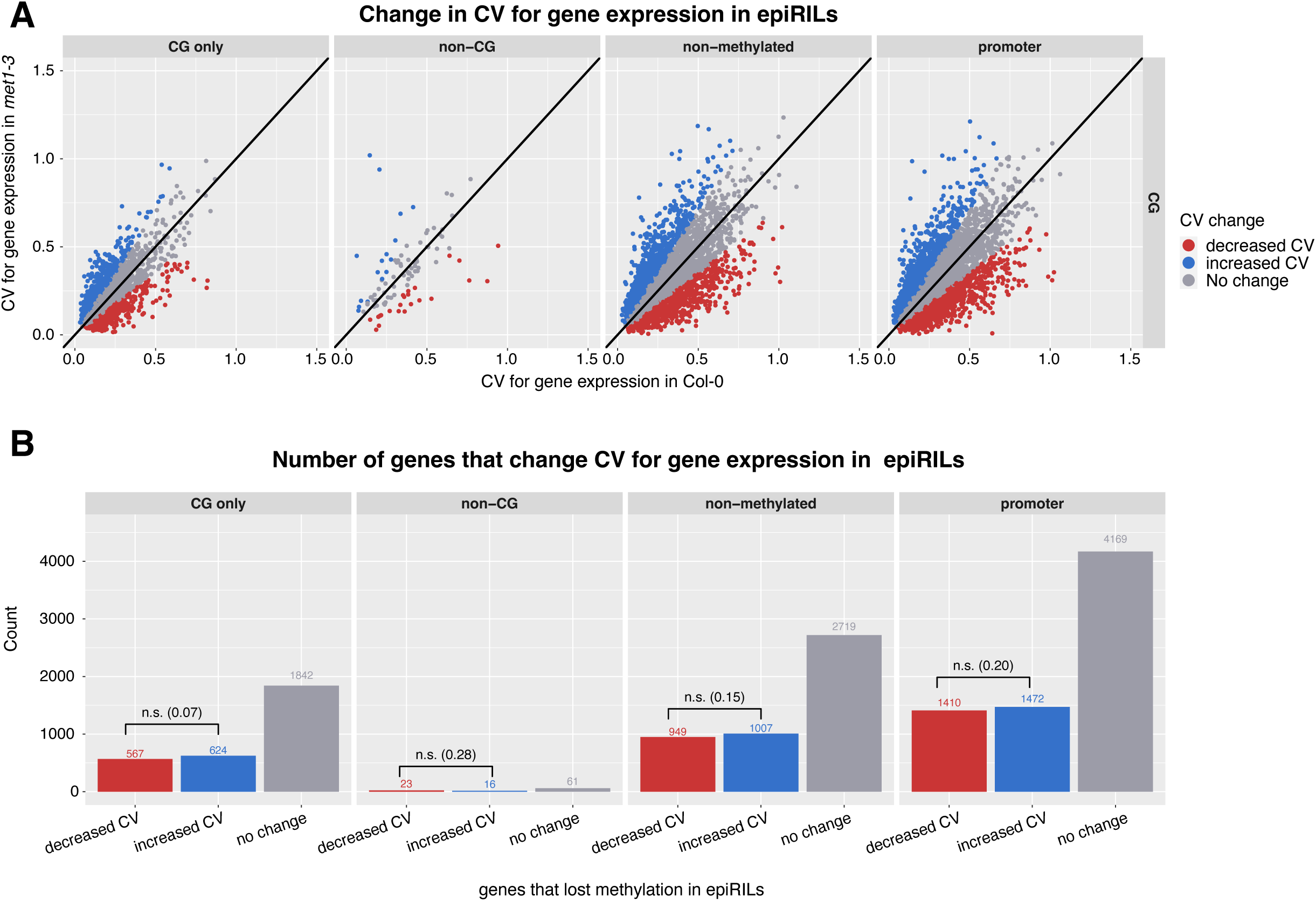
Changes in noise in gene expression in ten epiRIL lines. *(A)* Comparison between the Coefficient of Variation (CV) in genes in Col-0 background and genes in *met1-3* background. We considered in this analysis only genes without significant change in gene expression when in Col-0 and the mutant backgrounds but appearing in *met1-3* background in at least two epiRIL lines. Genes are grouped based on their coefficient of variation change: (blue) increased for genes with CV fold change greater than 0.5, (red) decreased for genes with CV fold change less than –0.5, and (grey) no change for those in-between. *(B)* Number of non-differentially expressed genes with loss of DNA methylation between Col-0 and *met1-3* backgrounds in the different categories shown in panel A.

**Supplementary Figure S12.**
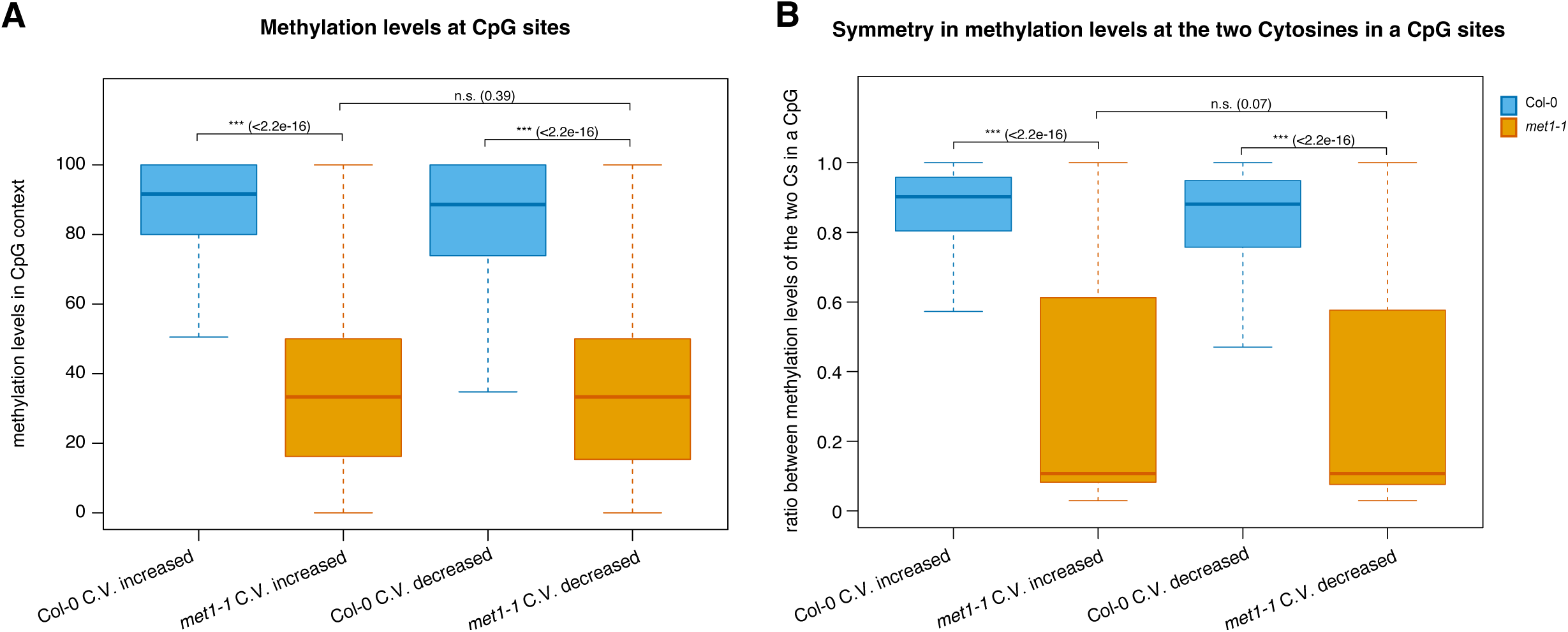
met1-1 mutant displays asymmetry of CpG methylation. *(A)* The methylation levels of all two cytosines in CpG context within gene bodies. We considered all CpGs that have at least 25% methylation for at least one of the two cytosines in Col-0 plants. We then subset those that overlapped genes that displayed either increase or decrease in variability in gene expression in *met1-1* mutant, while maintaining the same expression levels as in Col-0 (the CpG only in Figure 3B). We also performed the Mann–Whitney U test with corresponding P-value added to the plot. *(B)* Boxplot represents the proportion of CpG cytosine pairs with symmetric methylation (< 20% difference between the two cytosines) and the total number of CpG site for which at least one cytosine is 25% methylated in Col-0 and *met1-1*. We also performed the Mann–Whitney U test between the symmetry ratio in Col-0 and *met1-1* and between CpGs in genes that display increase variability and CpGs in genes that display decrease variability in *met1-1* (corresponding P-value added to the plot).

**Supplementary Figure S13.**
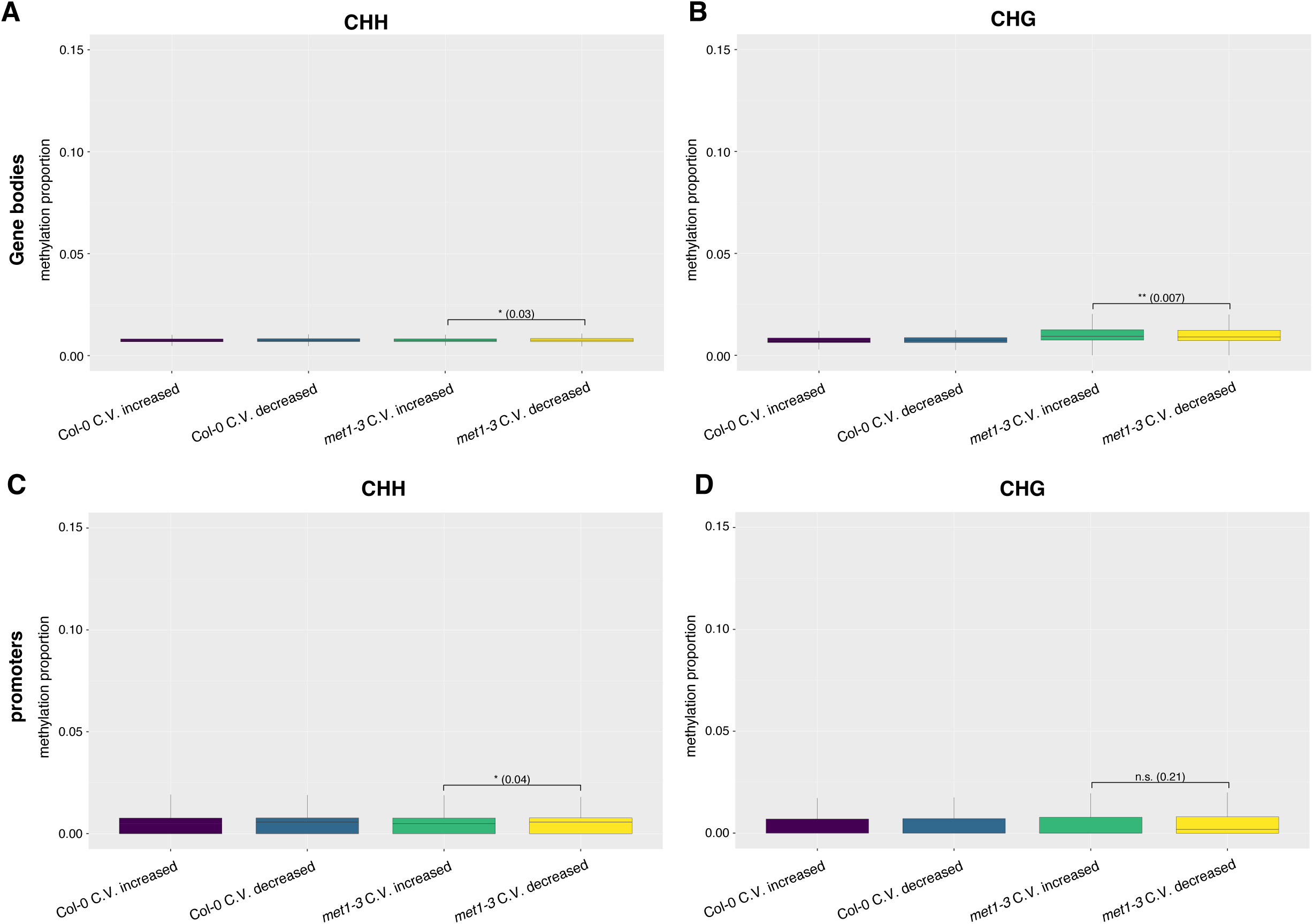
non-CpG methylation in met1-1 mutant at genes that display increase or decrease of noise in transcription. Methylation level of cytosines in (*A* and *C*) CHH and (*B* and *D*) CHG context located (*A-B*) within genes or (*C-D*) promoters of genes that displayed an increase or decrease in variability in gene expression in *met1-1* while maintaining the same expression levels as in Col-0. We performed the Mann–Whitney U test with corresponding P-value added to the plots.

**Supplementary Figure S14.**
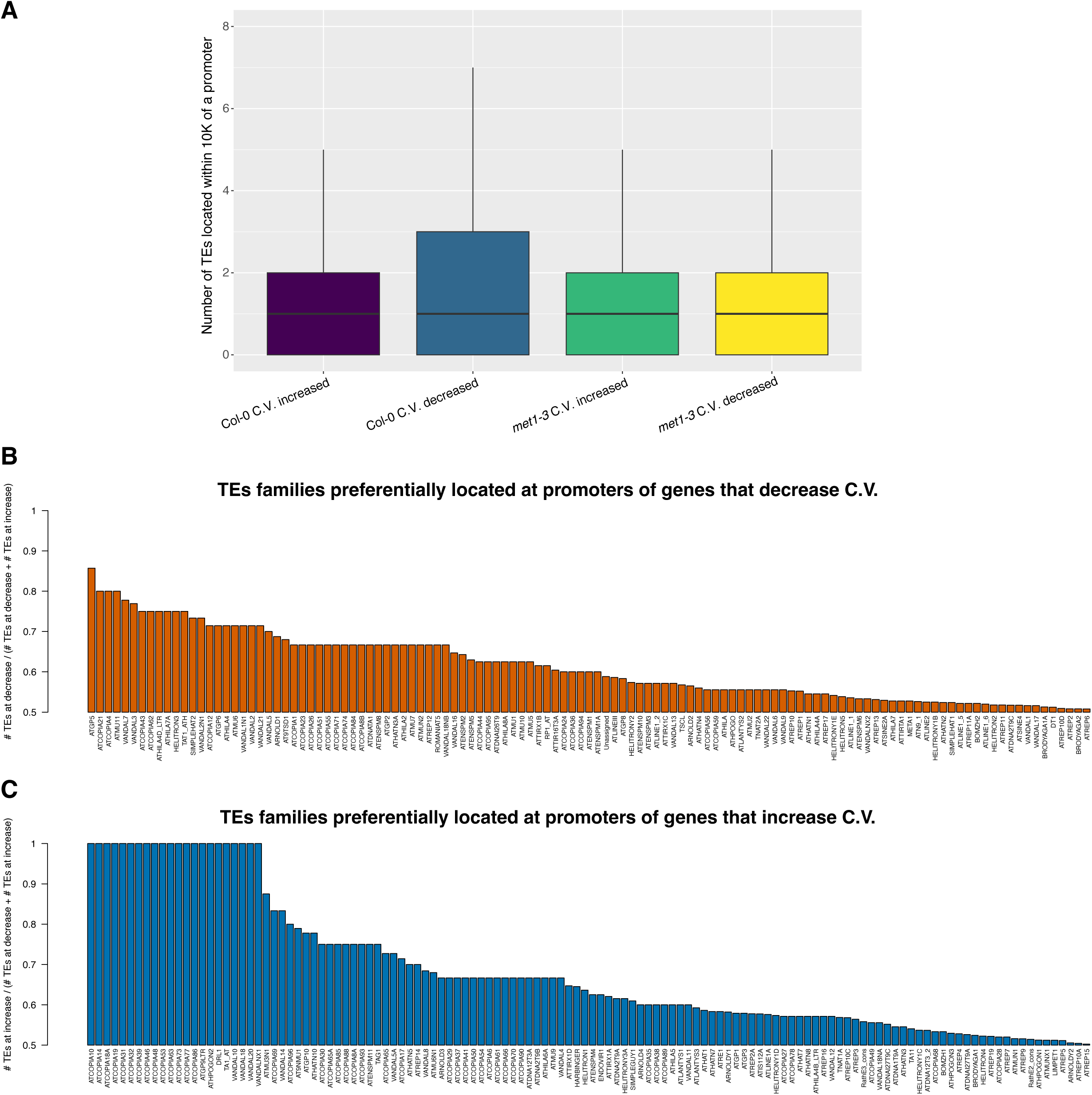
Transposable elements near promoters of genes that display increase or decrease of noise in transcription. *(A)* Number of TEs near promoters of genes that displayed an increase or decrease in variability in gene expression in *met1-3* while maintaining the same expression levels as in Col-0. *(B)* Barplot representing the number of copies of different TEs located within 10Kb of a promoter of a gene that displays lower noise in *met1-3*. The plot presents the TE families that have a higher proportion of copies near promoters of genes that display lower noise in *met1-3* (ratio between number of copies near genes that decrease C.V. and number of copies at both genes that increase and decrease C.V.)*. (C)* Barplot representing the number of copies of different TEs located within 10Kb of a promoter of a gene that displays higher noise in *met1-3*. The plot presents the TE families that have a higher proportion of copies near promoters of genes that display higher noise in *met1-3* (ratio between number of copies near genes that increase C.V. and number of copies at both genes that increase and decrease C.V.).

**Supplementary Table S1.** List of genes and the change in gene expression variability and methylation in *met1-1* mutant.

**Supplementary Table S2.** List of genes that have methylated promoter and the change in gene expression variability and methylation in *met1-1* mutant.

**Supplementary Table S3.** List of genes and the change in gene expression variability and methylation in *met1-3* mutant.

**Supplementary Table S4.** List of genes that have methylated promoter and the change in gene expression variability and methylation in *met1-3* mutant.

**Supplementary Table S5.** The epigenetic state and gene expression levels of all genes in epiRIL lines.

**Supplementary Table S6.** List of genes and the change in gene expression variability and methylation in epiRILs.

**Supplementary Table S7.** List of genes that have methylated promoter and the change in gene expression variability and methylation in epiRILs.

## Acknowledgements

The authors acknowledge the use of the High Performance Computing Facility at the University of Essex and would like to thank Stuart Newman for his support.

## Author contributions

M.C. U.B. and N.R.Z. conceived and designed the project. J.Z., R.A.E. and L.A.M. performed the analysis. M.C. performed the experiments. M.C., U.B. and N.R.Z. supervised the work. J.Z., M.C., U.B. and N.R.Z. wrote the paper. The authors read and approved the final manuscript.

## References

Abley K, Formosa-Jordan P, Tavares H, Chan EYT, Afsharinafar M, Leyser O & Locke JCW (2021) An aba-ga bistable switch can account for natural variation in the variability of arabidopsis seed germination time. Elife 10

Abley K, Goswami R & Locke JCW (2024) Bet-hedging and variability in plant development: Seed germination and beyond. Philosophical Transactions of the Royal Society B: Biological Sciences 379 doi:10.1098/rstb.2023.0048 [PREPRINT]

Abley K & Locke J (2021) Noisy transcription under the spotlight. Nat Plants 7: 1–2

Alamos S, Reimer A, Niyogi KK & Garcia HG (2021) Quantitative imaging of RNA polymerase II activity in plants reveals the single-cell basis of tissue-wide transcriptional dynamics. Nat Plants 7: 1037–1049

Alvarez-Fernandez R, Penfold CA, Galvez-Valdivieso G, Exposito-Rodriguez M, Stallard EJ, Bowden L, Moore JD, Mead A, Davey PA, Matthews JSA, et al (2021a) Time-series transcriptomics reveals a BBX32-directed control of acclimation to high light in mature Arabidopsis leaves. The Plant Journal 107: 1363–1386

Alvarez-Fernandez R, Penfold CA, Galvez-Valdivieso G, Exposito-Rodriguez M, Stallard EJ, Bowden L, Moore JD, Mead A, Davey PA, Matthews JSA, et al (2021b) Time-series transcriptomics reveals a BBX32-directed control of acclimation to high light in mature Arabidopsis leaves. Plant Journal 107: 1363–1386

Anders S & Huber W (2010) Differential expression analysis for sequence count data. Genome Biol 11

Anders S, Pyl PT & Huber W (2015) HTSeq-A Python framework to work with high-throughput sequencing data. Bioinformatics 31: 166–169

Araújo IS, Pietsch JM, Keizer EM, Greese B, Balkunde R, Fleck C & Hülskamp M (2017) Stochastic gene expression in Arabidopsis thaliana. Nat Commun 8

Arkin A, Ross J & Mcadams HH (1998) Stochastic Kinetic Analysis of Developmental Pathway Bifurcation in Phage-Infected Escherichia coli Cells

Bar-Even A, Paulsson J, Maheshri N, Carmi M, O’Shea E, Pilpel Y & Barkai N (2006) Noise in protein expression scales with natural protein abundance. Nat Genet 38: 636–643

Bechtold U, Penfold CA, Jenkins DJ, Legaie R, Moore JD, Lawson T, Matthews JSA, Vialet-Chabrand SRM, Baxter L, Subramaniam S, et al (2016) Time-series transcriptomics reveals that AGAMOUS-LIKE22 affects primary metabolism and developmental processes in drought-stressed arabidopsis. Plant Cell 28: 345–366

Berardini TZ, Mundodi S, Reiser L, Huala E, Garcia-Hernandez M, Zhang P, Mueller LA, Yoon J, Doyle A, Lander G, et al (2004) Functional annotation of the Arabidopsis genome using controlled vocabularies. Plant Physiol 135: 745–755

Bewick AJ & Schmitz RJ (2017) Gene body DNA methylation in plants. Curr Opin Plant Biol 36: 103– 110 doi:10.1016/j.pbi.2016.12.007 [PREPRINT]

Bishop AL, Rab FA, Sumner ER & Avery S V. (2007) Phenotypic heterogeneity can enhance rare-cell survival in ‘stress-sensitive’ yeast populations. Mol Microbiol 63: 507–520

Bolger AM, Lohse M & Usadel B (2014) Trimmomatic: A flexible trimmer for Illumina sequence data. Bioinformatics 30: 2114–2120

Bolstad BM, Irizarry RA, Astrand M° & Speed TP (2003) A comparison of normalization methods for high density oligonucleotide array data based on variance and bias

Brettschneider J, Collin F, Bolstad BM & Speed TP (2007) Quality assessment for short oligonucleotide microarray data.

Catoni M & Cortijo S Running title: EpiRILs: lessons from Arabidopsis EpiRILs: lessons from Arabidopsis

Catoni M, Griffiths J, Becker C, Zabet NR, Bayon C, Dapp M, Lieberman-Lazarovich M, Weigel D & Paszkowski J (2017) DNA sequence properties that predict susceptibility to epiallelic switching. EMBO J 36: 617–628

Catoni M, Tsang JMF, Greco AP & Zabet NR (2018) DMRcaller: A versatile R/Bioconductor package for detection and visualization of differentially methylated regions in CpG and non-CpG contexts. Nucleic Acids Res 46

Catoni M & Zabet NR (2021) Analysis of Plant DNA Methylation Profiles Using R. In Plant Transposable Elements: Methods and Protocols, Cho J (ed) pp 219–238. New York, NY: Springer US

Cortijo S, Aydin Z, Ahnert S & Locke JC (2019) Widespread inter-individual gene expression variability in Arabidopsis thaliana. Mol Syst Biol 15

Eldar A & Elowitz MB (2010) Functional roles for noise in genetic circuits. Nature 467: 167–173 doi:10.1038/nature09326 [PREPRINT]

Elowitz MB, Levine AJ, Siggia ED & Swain PS (2002) Stochastic Gene Expression in a Single Cell. Science (1979) 297: 1183–1186

Erdmann RM & Picard CL (2020) RNA-directed DNA Methylation. PLoS Genet 16: e1009034

Folta A, Severing EI, Krauskopf J, van de Geest H, Verver J, Nap JP & Mlynarova L (2014) Over-expression of Arabidopsis AtCHR23 chromatin remodeling ATPase results in increased variability of growth and gene expression. BMC Plant Biol 14

Fultz D, Choudury SG & Slotkin RK (2015) Silencing of active transposable elements in plants. Curr Opin Plant Biol 27: 67–76 doi:10.1016/j.pbi.2015.05.027 [PREPRINT]

Gygi SP, Rochon Y, Franza BR & Aebersold R (1999) Correlation between Protein and mRNA Abundance in Yeast. Mol Cell Biol 19: 1720–1730

Hagai T, Chen X, Miragaia RJ, Rostom R, Gomes T, Kunowska N, Henriksson J, Park JE, Proserpio V, Donati G, et al (2018) Gene expression variability across cells and species shapes innate immunity. Nature 563: 197–202

Hani S, Cuyas L, David P, Secco D, Whelan J, Thibaud MC, Merret R, Mueller F, Pochon N, Javot H, et al (2021) Live single-cell transcriptional dynamics via RNA labelling during the phosphate response in plants. Nat Plants 7: 1050–1064

Horvath R, Laenen B, Takuno S & Slotte T (2019) Single-cell expression noise and gene-body methylation in Arabidopsis thaliana. Heredity (Edinb) 123: 81–91

Huang DW, Sherman BT & Lempicki RA (2009) Systematic and integrative analysis of large gene lists using DAVID bioinformatics resources. Nat Protoc 4: 44–57

Huh I, Zeng J, Park T & Yi S V (2013) DNA methylation and transcriptional noise

Inagaki S, Takahashi M, Hosaka A, Ito T, Toyoda A, Fujiyama A, Tarutani Y & Kakutani T (2017) Gene-body chromatin modification dynamics mediate epigenome differentiation in *Arabidopsis*. EMBO J 36: 970–980

Kim D, Pertea G, Trapnell C, Pimentel H, Kelley R & Salzberg SL (2013) TopHat2: Accurate alignment of transcriptomes in the presence of insertions, deletions and gene fusions. Genome Biol 14

Krueger F & Andrews SR (2011) Bismark: A flexible aligner and methylation caller for Bisulfite-Seq applications. Bioinformatics 27: 1571–1572

Liang Z, Shen L, Cui X, Bao S, Geng Y, Yu G, Liang F, Xie S, Lu T, Gu X, et al (2018) DNA N6-Adenine Methylation in Arabidopsis thaliana. Dev Cell 45: 406–416.e3

Lister R, O’Malley RC, Tonti-Filippini J, Gregory BD, Berry CC, Millar AH & Ecker JR (2008) Highly Integrated Single-Base Resolution Maps of the Epigenome in Arabidopsis. Cell 133: 523–536

Liu N, Fromm M & Avramova Z (2014) H3K27me3 and H3K4me3 chromatin environment at super-induced dehydration stress memory genes of arabidopsis thaliana. Mol Plant 7: 502–513

Maintainer SG & Grote S (2024) Type Package Title Gene ontology enrichment using FUNC

Matzke MA, Kanno T & Matzke AJM (2015) RNA-directed DNA methylation: The evolution of a complex epigenetic pathway in flowering plants. Annu Rev Plant Biol 66: 243–267

Mi H, Dong Q, Muruganujan A, Gaudet P, Lewis S & Thomas PD (2009) PANTHER version 7: Improved phylogenetic trees, orthologs and collaboration with the Gene Ontology Consortium. Nucleic Acids Res 38

Neri F, Rapelli S, Krepelova A, Incarnato D, Parlato C, Basile G, Maldotti M, Anselmi F & Oliviero S (2017) Intragenic DNA methylation prevents spurious transcription initiation. Nature 543: 72–77

Reinders J, Wulff BBH, Mirouze M, Marí-Ordóñez A, Dapp M, Rozhon W, Bucher E, Theiler G & Paszkowski J (2009) Compromised stability of DNA methylation and transposon immobilization in mosaic Arabidopsis epigenomes. Genes Dev 23: 939–950

Rigal M, Kevei Z, Pélissier T & Mathieu O (2012) DNA methylation in an intron of the IBM1 histone demethylase gene stabilizes chromatin modification patterns. EMBO J 31: 2981–2993

Saze H, Shiraishi A, Miura A & Kakutani T (2008) Control of Genic DNA Methylation by a jmjC Domain-Containing Protein in Arabidopsis thaliana. Science (1979) 319: 462–465

Schoech AP & Zabet NR (2014) Facilitated diffusion buffers noise in gene expression. Phys Rev E Stat Nonlin Soft Matter Phys 90

Sclep G, Allemeersch J, Liechti R, De Meyer B, Beynon J, Bhalerao R, Moreau Y, Nietfeld W, Renou JP, Reymond P, et al (2007) CATMA, a comprehensive genome-scale resource for silencing and transcript profiling of Arabidopsis genes. BMC Bioinformatics 8

Spudich JL & Koshland DE (1976) Non-genetic individuality: chance in the single cell

Stroud H, Greenberg MVC, Feng S, Bernatavichute Y V. & Jacobsen SE (2013) Comprehensive analysis of silencing mutants reveals complex regulation of the Arabidopsis methylome. Cell 152: 352–364

Stroud H, Hale CJ, Feng S, Caro E, Jacob Y, Michaels SD & Jacobsen SE (2012) DNA methyltransferases are required to induce heterochromatic re-replication in arabidopsis. PLoS Genet 8

Volfson D, Marciniak J, Blake WJ, Ostroff N, Tsimring LS & Hasty J (2006) Origins of extrinsic variability in eukaryotic gene expression. Nature 439: 861–864

Weinberger LS, Burnett JC, Toettcher JE, Arkin AP & Schaffer D V. (2005) Stochastic gene expression in a lentiviral positive-feedback loop: HIV-1 Tat fluctuations drive phenotypic diversity. Cell 122: 169–182

Wu S, Li K, Li Y, Zhao T, Li T, Yang YF & Qian W (2017) Independent regulation of gene expression level and noise by histone modifications. PLoS Comput Biol 13

Zabet NR (2011) Negative feedback and physical limits of genes. J Theor Biol 284: 82–91

Zabet NR, Catoni M, Prischi F & Paszkowski J (2017a) Cytosine methylation at CpCpG sites triggers accumulation of non-CpG methylation in gene bodies. Nucleic Acids Res 45

Zabet NR, Catoni M, Prischi F & Paszkowski J (2017b) Cytosine methylation at CpCpG sites triggers accumulation of non-CpG methylation in gene bodies. Nucleic Acids Res 45: 3777–3784

Zabet NR & Chu DF (2010) Computational limits to binary genes. J R Soc Interface 7: 945–954

Zilberman D, Coleman-Derr D, Ballinger T & Henikoff S (2008) Histone H2A.Z and DNA methylation are mutually antagonistic chromatin marks. Nature 456: 125–129

